# PKD2 is an essential ion channel subunit in the primary cilium of the renal collecting duct epithelium

**DOI:** 10.1101/215814

**Authors:** Xiaowen Liu, Thuy Vien, Jingjing Duan, Shu-Hsien Sheu, Paul G. DeCaen, David E. Clapham

**Affiliations:** Howard Hughes Medical Institute, Department of Cardiology, Boston Children's Hospital, 320 Longwood Avenue, Boston, MA 02115, USA; Department of Neurobiology, Harvard Medical School, Boston, MA 02115, USA; Northwestern University, Feinberg School of Medicine, Department of Pharmacology, 320 East Superior Street, Chicago, Illinois 60611

## Abstract

Mutations in either *Pkd1* or *Pkd2* result in Autosomal Dominant Polycystic Kidney Disease (ADPKD). Although PKD2 is proposed to be an ion channel subunit, recordings of PKD2 ion channels conflict in their properties. Using a new ADPKD mouse model, we observe primary cilia are abnormally long in cells associated with cysts. Using primary cultures of collecting duct epithelial cells, we show that PKD2, but not PKD1, is a required subunit for primary cilia ion channel. The ciliary PKD2 channel conducts potassium and sodium ions, but little calcium. We also demonstrate that PKD2 is not constitutively active in the plasma membrane, but PKD2 channels are functional in primary cilia and are sensitized by high cilioplasmic [Ca^2+^]. We introduce a novel method for measuring PKD2 channels heterologously expressed in primary cilia of HEK-293 cells, which will have utility characterizing *Pkd2* variants that cause ADPKD in their native ciliary membrane.

## INTRODUCTION

Autosomal dominant polycystic kidney disease (ADPKD) is an adult-onset disease characterized by focal cyst development resulting from heterozygous mutations in *Pkd1* or *Pkd2*(Brasier & Henske, 1997; Grantham, 2001; Hughes et al., 1995; Mochizuki et al., 1996). While considered a dominant monogenic disease, the prevailing two-hit model states that ADPKD is recessive at the cellular level and that cysts develop from cells after acquiring a second somatic mutation to deactivate the remaining normal allele(Koptides, Hadjimichael, Koupepidou, Pierides, & Constantinou Deltas, 1999; Pei, 2001; F. Qian et al., 1997; Wu et al., 1998). Mouse models of ADPKD implicate ciliary PKD1 and PKD2 dysfunction in kidney cyst formation. Complete genetic knockout of either *Pkd1* or *Pkd2* in mice results in embryonic lethality due to structural defects in the cardiovascular system, pancreas, and kidneys(Boulter et al., 2001; K. Kim, Drummond, Ibraghimov-Beskrovnaya, Klinger, & Arnaout, 2000; Lu et al., 1997; Wu & Somlo, 2000; Wu et al., 2002). The onset of kidney cyst development in adult mice following conditional inactivation of *Pkd1* or the intraflagellar transport protein kinesin, KIF3a (required for cilia formation), progresses well into adulthood, in analogy to the late progression of ADPKD in humans(Davenport et al., 2007; Piontek, Menezes, Garcia-Gonzalez, Huso, & Germino, 2007; Shibazaki et al., 2008). Kidney-specific conditional repression of either *Pkd1* (*Pax8^rtTA^*; *TetO-cre*; *Pkd1^fl/fl^*) (Shibazaki et al., 2008)or *Pkd2* (*Pax8^rtTA^; TetO-cre; Pkd2^fl/fl^*)(M. Ma, Tian, Igarashi, Pazour, & Somlo, 2013) develop cysts within 10 weeks after the start of doxycycline induction, suggesting that expression of both genes is necessary to prohibit cyst development in mature mice. Recently, the cystic phenotype found in mice deficient in *Pkd2* can be dose-dependently rescued by *Pkd2* transgene expression(A. Li et al., 2015). Our poor understanding of the functional properties of PKD1, PKD2, and the PKD1/PKD2 complex impedes the development of pharmacological therapeutic strategies - ADPKD is currently treated by dialysis and kidney transplant(LaRiviere, Irazabal, & Torres, 2015). Since PKD2’s function is unclear, it is not currently known if all ADPKD-causing variants in *Pkd2* cause a loss of function of the putative ion channel in primary cilia.

PKD1 (PC1) is predicted to adopt an 11-transmembrane topology with a large autocleaved (via a G protein-coupled receptor proteolytic site GPS) amino-terminal ectodomain (>3,000 residues)(Harris, Ward, Peral, & Hughes, 1995) that is comprised of an array of putative adhesion and ligand-binding modules(Burn et al., 1995; Hughes et al., 1995; F. Qian et al., 2002). PKD2 (or PC2, TRPP1, formerly TRPP2) is a member of the large, 6-transmembrane spanning transient receptor potential (TRP) ion channel family(Ramsey, Delling, & Clapham, 2006; Venkatachalam & Montell, 2007) and has been observed to form a complex with PKD1. It is proposed to interact with PKD2 through a probable coiled-coil domain(Newby et al., 2002; F. Qian et al., 1997; Tsiokas, Kim, Arnould, Sukhatme, & Walz, 1997), but in other experiments, the PKD1 and PKD2 interaction is preserved in overexpressed systems without the coiled-coil domain and is dependent on the N-terminal domain(Babich et al., 2004; Celic et al., 2012; Feng et al., 2008). Based on biochemistry and immunoreactivity, both proteins can be found in the primary cilium and ER(Ong & Wheatley, 2003; Yoder, Hou, & Guay-Woodford, 2002). In addition, some studies suggest that PKD1 and PKD2 may reciprocally affect each other’s surface membrane or ciliary localization(Harris et al., 1995; Ong & Wheatley, 2003; C. Xu et al., 2007). A recent study using inner medullary collecting duct (IMCD) cells derived from human ADPKD cysts suggests that impairing the function of PKD1 or PKD2 negatively affects the localization of the other protein: cells expressing an ADPKD-associated PKD1 mutation that prevents GPS domain cleavage have decreased amounts of both PKD1 and PKD2 in their primary cilia(C. Xu et al., 2007). As discussed below, our understanding of ADPKD pathogenesis is hampered by disagreements about the basic properties of the putative PKD2 current. Furthermore, little is understood regarding what role (if any) the primary cilia have in controlling the progression of cyst formation in ADPKD. Nonetheless, there is no ambiguity in the finding that mutations in *Pkd1* or *Pkd2* are genetically linked to formation of cysts in kidney and other tissues to cause significant morbidity and mortality in humans(Mochizuki et al., 1996; Ong & Harris, 2015)

Previous work reported single channel events in artificial bilayers attributable to an ion channel complex involving PKD1 and PKD2(Delmas et al., 2004; Gonzalez-Perrett et al., 2001; Hanaoka et al., 2000). However, exogenous expression or reconstitution of PKD2 in artificial bilayers have produced contradictory ion selectivity and voltage dependence. The PKD2 ion channel was initially reported to conduct calcium(Gonzalez-Perrett et al., 2001; Hanaoka et al., 2000), and was also blocked by calcium(Cai et al., 2004). Most recently, a gain-of-function mutation (F604P), but not *wt* PKD2, underlies a measurable current when heterologously expressed in *Xenopus* oocytes(Arif Pavel et al., 2016). This study demonstrated that plasma membrane PKD2 expression does not appear to be hampered by the lack of PKD1, but rather that native PKD2 channels appears to be constitutively closed unless mutated (F604P). The monovalent-selective current of this mutant is blocked by divalent ions (Ca^2+^ and Mg^2+^). As we will review in the Discussion, several putative PKD2 activators have been reported to sensitize PKD2 channels, but we have been unable to reproduce these results(S. Kim et al., 2016; Leuenroth et al., 2007). While the apparent differences observed can be rooted in methodology, these preparations are measured from non-native membranes and thus share the same disadvantage. Recently, two methods have been used to measure ion currents from intact primary cilia(DeCaen, Delling, Vien, & Clapham, 2013; N. K. Kleene & Kleene, 2012), thus preserving their unique native microenvironment without the need for reconstitution.

For this study, we crossed our *Arl13B-EGFP^tg^* strain(DeCaen et al., 2013) with *Pax8^rtTA^; TetOcre*; *Pkd1^fl/fl^* (*cPkd1*) or *Pax8^rtTA^; TetO-cre*; *Pkd2^fl/fl^* (*cPkd2*) mice provided by the Somlo lab(M. Ma et al., 2013; Shibazaki et al., 2008). The progeny express the Arl13B-EGFP^tg^ cilia reporter, and ablation of either *Pkd1* or *Pkd2* genes expression in the kidney under the Pax8^rtTA^ promotor(Traykova-Brauch et al., 2008) is *TetO-cre* doxycycline-dependent. For brevity, we will call these animal strains either *Arl13B-EGFP^tg^*:*cPkd1* or *Arl13B-EGFP^tg^*:*cPkd2*, respectively. Consistent with previous reports(M. Ma et al., 2013), we find that repression of either *Pkd1* or *Pkd2* results in obvious kidney cysts within two months after removal of doxycycline. Primary cilia from cysts of either doxycycline treated *Arl13B-EGFP^tg^:cPkd1* or *Arl13B-EGFP^tg^:cPkd2* mice were substantially elongated compared to control littermates. We utilized the cilia patch method to directly measure putative ion channels from primary cultures of inner medullary collecting duct epithelial cells (pIMCD) from these mice. We characterize the native ciliary PKD2 currents, which can be conditionally ablated using doxycycline in the *Arl13B-EGFP^tg^*:*cPkd2* mouse model.

Surprisingly, PKD2 forms a functional ion channel in primary cilia without PKD1 expression, calling into question the hypothesis that PKD1 is an obligate subunit of putative PKD1/PKD2 heteromeric ion channels. The ciliary PKD2 current preferentially conducts the monovalents K^+^ and Na^+^, over divalent Ca^2+^ ions. Millimolar external [Ca^2+^] weakly permeates through the PKD2 pore and blocks the inward sodium current. The open probability of PKD2 is enhanced by internal calcium (EC_50_ = 1.3 μM), slightly exceeding the resting cilioplasmic [Ca^2+^] (~300-600 nM)(Delling, DeCaen, Doerner, Febvay, & Clapham, 2013; Delling et al., 2016). Native constitutive plasma membrane currents are not affected by conditional ablation of either *Pkd1* or *Pkd2* from pIMCD cells. Thus, we find no evidence for homomeric or heteromeric polycystin channels in the plasma membrane. Heterologous, stably-expressed PKD2-GFP traffics to the primary cilia of HEK-293 cells, where cilia patch clamp recordings recapitulate the ion selectivity and internal calcium potentiation effects observed in primary cilia of native pIMCD cells.

## RESULTS

### Progressive cyst formation in a new mouse model

Previous work demonstrated that the human ADPKD kidney cyst phenotype can be reproduced in mice 14 weeks after conditional ablation of nephron-localized *Pkd1* or *Pkd2*(M. Ma et al., 2013). To understand the putative ciliary ion channel function of PKD1 or/and PKD2 and to determine the effects of kidney cyst formation on cilia morphology, we crossed our *Arl13B-EGFP^tg^* strain(DeCaen et al., 2013) with *cPkd1* or *cPkd2* mice (provided by the Somlo lab). We then induced either *Pkd1* or *Pkd2* gene inactivation in adult animals (~P28) by introducing doxycycline (2 mg/ml or 3.9 mM) into the drinking water for two weeks. After this treatment period, doxycycline was removed and kidney histology was performed from 2 and 4-month post-treatment animals (**Figure 1A**, **Figure 1-figure supplement 1**, **Figure 1-figure supplement 2A**).

**Figure 1.**
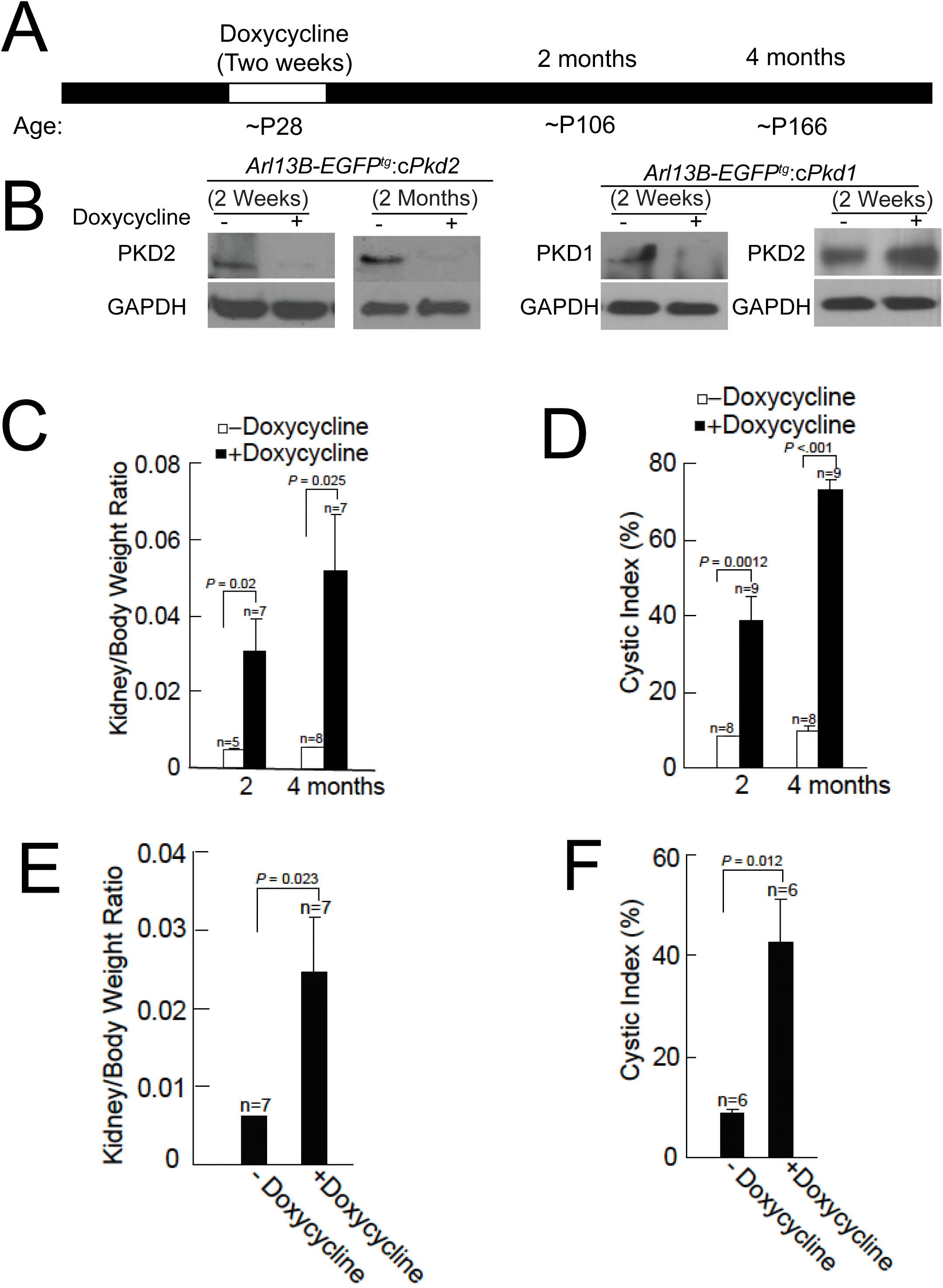
Onset of kidney tubule cyst formation in *Arl13B-EGFP^tg^:cPkd1* and *Arl13B-EGFP^tg^:cPkd2* animals. A) Study design to assess cyst formation after genetic ablation of either *Pkd1* or *Pkd2*. One protocol is shown that assesses PKD1 or PKD2 at P28 in the conditional knockout mouse. B) Loss of PKD1 and PKD2 protein expression as assessed by immunoblot, 2 weeks and 2 months after doxycycline removal. Whole cell lysates were prepared from pIMCD cells and subjected to western blot analysis. Three independent experiments were performed. C) Kidney weight / body weight was increased in *Arl13B-EGFP^tg^:cPkd2* mice with doxycycline treatment compared to control littermates without doxycycline treatment. D) Cystic index (Methods) shows that cysts increased in *Arl13B-EGFP^tg^:cPkd2* mice. E) Kidney weight / body weight was increased in *Arl13B-EGFP^tg^:cPkd1* with doxycycline treatment (2 months after doxycycline removal) compared to control littermates without doxycycline treatment. F) Cystic index shows increased size and number of cysts in *Arl13B-EGFP^tg^:cPkd1* mice

Immunoblots were performed from pIMCD cell lysates from 2-week and 2-month doxycycline-ablated *Arl13B-EGFP^tg^*:*cPkd2* or 2-week post-treatment of *Arl13B-EGFP^tg^*:*cPkd1* mice, indicating that the recombinase stably abolished PKD2 or PKD1 protein expression (**Figure 1B**). PKD2 expression from *Arl13B-EGFP^tg^:cPkd1* animals was unaffected by PKD1 ablation (**Figure 1B**). Consistent with previous reports from the *cPkd2 and cPkd1* strain, we observed kidney cyst formation in *Arl13B-EGFP^tg^:cPkd2* and *Arl13B-EGFP^tg^:cPkd1* mice (**Figure 1-figure supplement 1**, **Figure 1-figure supplement 2A**). The extent of cyst formation in these mice was quantified as the kidney-to-body weight ratio and cystic index (**Figure 1C-F**). Based on these measures, the cystic phenotype was progressive, as seen by comparing the post 2-month and 4-month treatment groups (**Figure 1C-F**, **Figure 1-figure supplement 1**, **Figure 1-figure supplement 2A**).

### Abnormal cilia in PKD1- or PKD2-ablated mice

Using confocal microscopy, we compared cilia morphology from kidneys of *Arl13B-EGFP^tg^:cPkd2* and *Arl13B-EGFP^tg^:cPkd1* mice treated with or without doxycycline (**Figure 2A-C**, **Figure 1-figure supplement 2B-D**). Here, we observed an ~3.2-fold increase in cilia length with the progression of ADPKD (5.7 ± 0.4 μm for 2 months and 18.4 ± 1.2 μm for 4 months post-treatment) with *Arl13B-EGFP^tg^:cPkd2* mice, whereas cilia length from control littermates did not differ substantially over the same time course (3.8 ± 0.16 μm and 4.5 ± 0.2 μm, respectively). Also, we found that cilia length from tubule cells lining cysts were ~4-times longer than from unaffected tubules from the same animals (**Figure 2C**) (12 ± 1.1 μm and 3.1 ± 0.2 μm, respectively). As for *Arl13B-EGFP^tg^:cPkd1* mice, we observed an ~2.4-fold increase in cilia length with the progression of ADPKD (4.1 ± 0.1 μm for control littermates and 9.9 ± 0.5 μm for 2 months post-treatment). These results demonstrate the neither PKD1 nor PKD2 expression is required for primary ciliogenesis from the tubule epithelium, but implies that PKD1 or PKD2 expression is somehow related to cilia length. Since aberrant cilia morphology was mostly found in cystic tissue epithelia compared to non-cystic tubules, ciliary PKD1 or PKD2 may regulate continuing renal tubular cell differentiation. However, it is unclear if irregular cilia morphology is a consequence or cause of cyst formation, and what function overexpression of Arl13B in combination with PKD1 or PKD2 ablation may have in maintaining normal cilia length.

**Figure 2.**
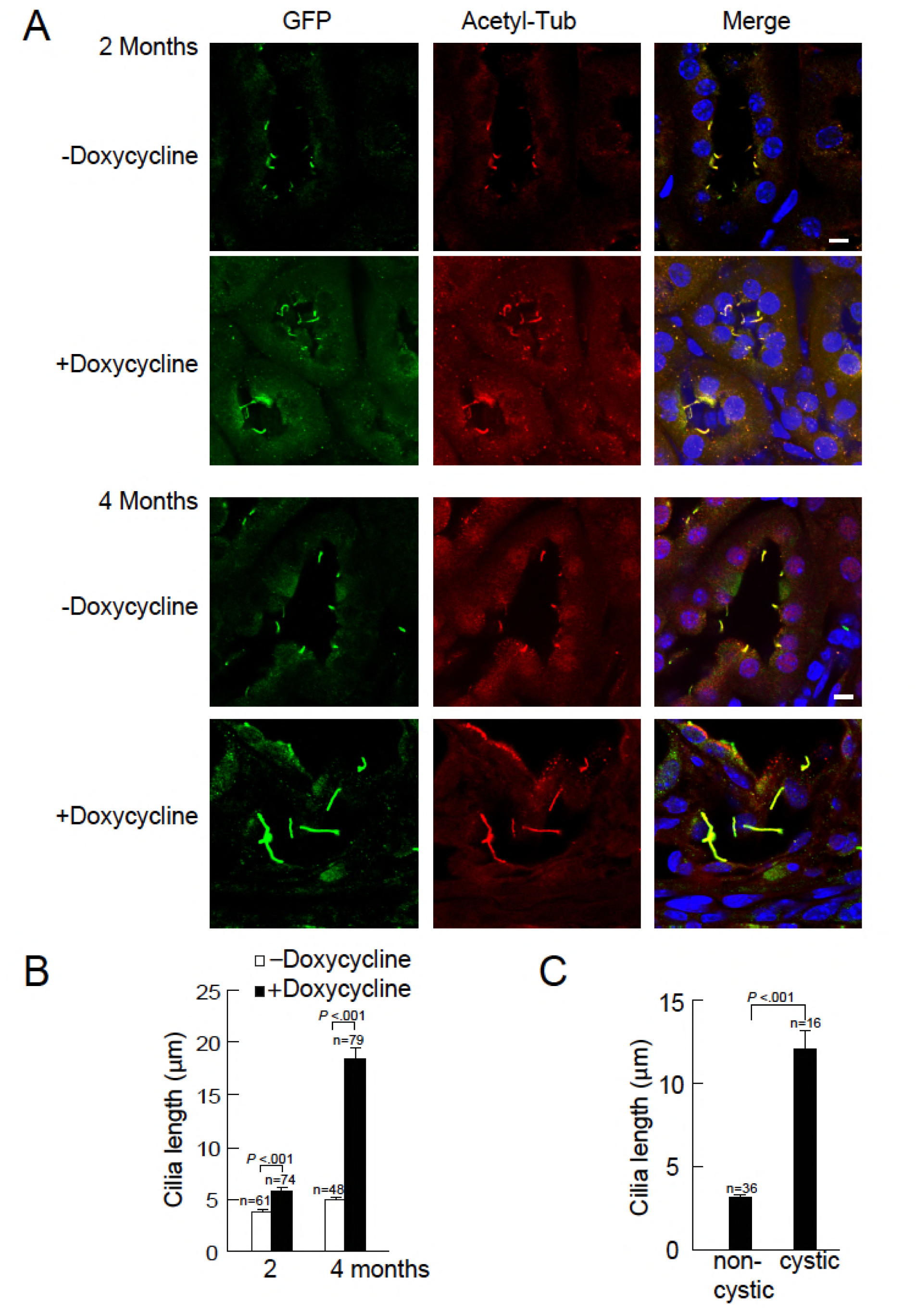
Cystic kidney cilia are abnormally long compared to unaffected tubules. A) Representative kidney sections from *Arl13B-EGFP^tg^:cPkd2* mice were immunolabeled with antibodies against EGFP and acetylated tubulin. Three independent experiments were performed. Scale bars, 5 μm. B) Cilia length was measured with the progression of cyst formation from kidney of Arl*13B-EGFP^tg^:cPkd2* mice. C) Cilia length was measured in cystic and non-cystic areas from the *Arl13B-EGFP^tg^:cPkd2* mice after 2 months of doxycycline removal.

### Ciliary trafficking and ion channel activity of PKD2 are independent of PKD1

Using animals from the same study design, we harvested pIMCD cells from *Arl13B-EGFP^tg^* mice. The cell membrane of the dissociated cells retained anti-aquaporin 2 antibody reactivity and Arl13B was found in the primary cilia of intact distal collecting ducts (**Figure 3A-C**). Using the validated antibody described in **Figure 1**, we confirmed the lack of ciliary PKD2 from cultured pIMCD cells from post-doxycycline-treated *Arl13B-EGFP^tg^*:*cPkd2* mice (**Figure 3D**). Importantly, the pIMCD cells isolated from post-doxycycline-treated *Arl13B-EGFP^tg^:cPkd1* animals retained their ciliary PKD2, suggesting that ciliary PKD2 trafficking does not require PKD1 (**Figure 3E**, **Figure 1-figure supplement 2D**).

**Figure 3.**
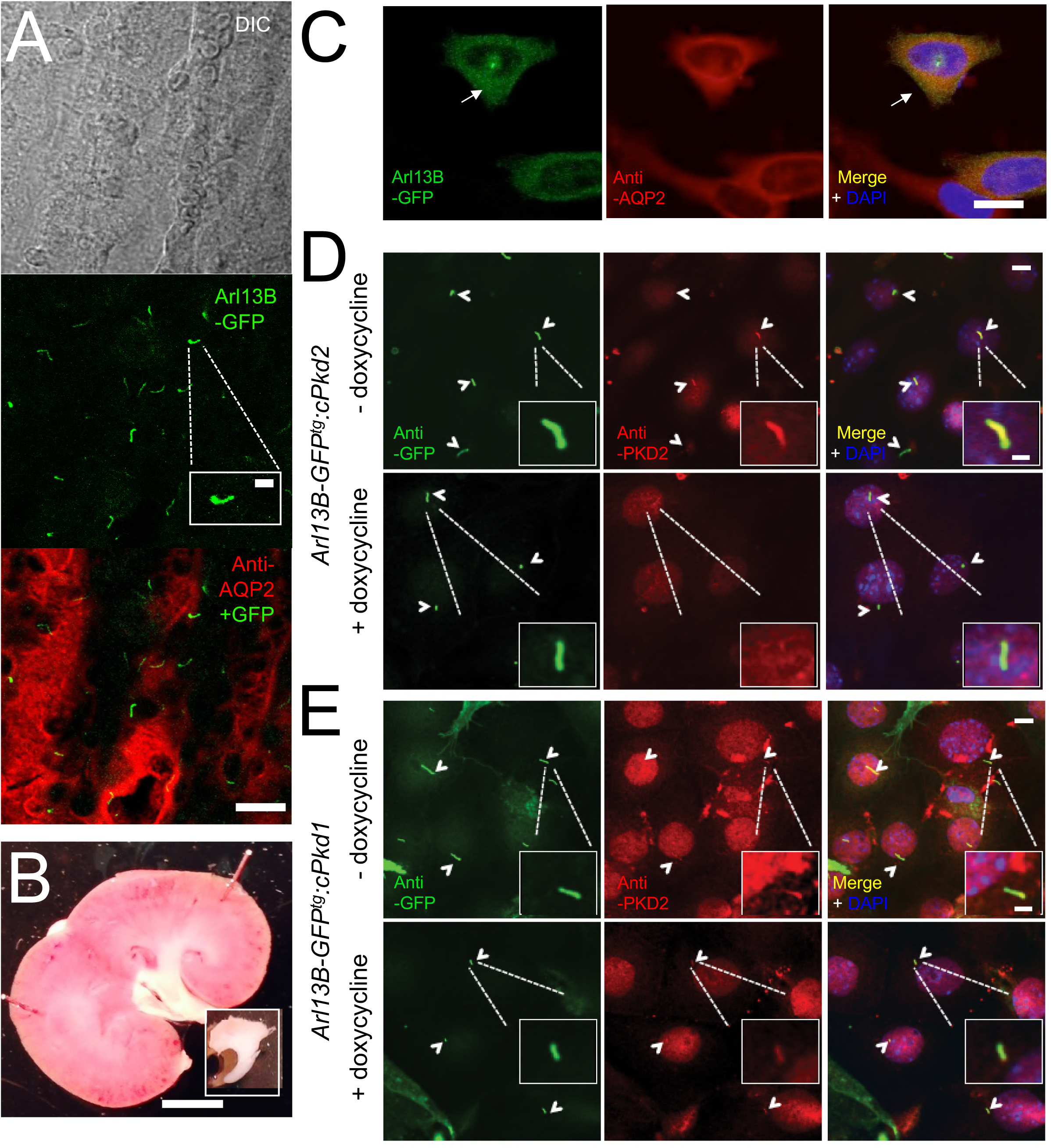
*In situ* and *in vitro* detection of ciliary PKD2 in *Arl13B-EGFP^tg^* pIMCD cells. A) Confocal images from a 100 μm-thick fixed kidney slice: DIC image in gray; Aquaporin 2 (kidney collecting duct epithelial cell epitope) labeled with Alexa-569 (red); Cilia *Arl13B-EGFP^tg^* (green). Three independent experiments were performed. Scale bar = 30 μm, inset image scale bar = 5 μm. B) Sagittal section of 3-month-old mouse kidney with the inner medulla removed (bottom inset). Three independent experiments were performed. Scale bar = 1 mm. C) Confocal images of fixed primary collecting duct epithelial cells after two days in culture, immunostained with anti-aquaporin 2 antibody in A). Three independent experiments were performed. Scale bar = 5 μm. D) Immunofluorescence using anti-GFP (green) and anti-PKD2 (red) showing the loss of PKD2 in pIMCD cells isolated from kidney papillae of *Arl13B-EGFP^tg^:cPkd2* mice. Three independent experiments were performed. 5 mice were used for each group. Arrowheads point to primary cilia. Scale bar = 5 μM, inset image scale bar = 1 μm. E) Immunofluorescence with anti-GFP (green) and anti-PKD2 (red), showing ciliary PKD2 in pIMCD cells isolated from kidney papillae of *Arl13B-EGFP^tg^:cPkd1* mice. Three independent experiments were performed. 5 mice were used for each group. Arrowheads point to primary cilia. Scale bar = 5 μM, inset image scale bar = 1 μm.

Next, we patch clamped pIMCD cells, in which the primary cilia could be visualized and expression of either PKD1 or PKD2 subunits of the putative cilia ion channel complex could be conditionally controlled. Previously, we used the cilia patch method to identify the heteromeric PKD1-L1/2-L1 channel in primary cilium of *Arl13B-EGFP^tg^* retinal pigmented epithelial cells (RPE) and mouse embryonic fibroblasts (MEF)(DeCaen et al., 2013). Also, we described, but did not identify, a large outward conductance channel (outward γ = 98 ± 2 pS) from the cilia of an immortalized IMCD-3 cell line, which has been characterized (outward γ = 96 pS) and subsequently identified as PKD2 by the Kleene group(S. J. Kleene & Kleene, 2017). Thus, we extended our cilia electrophysiology methods to test cilia ion currents from pIMCD cells and determine if PKD1 and/or PKD2 are subunits of the ion channel. After establishing high resistance seals (>16 GΩ) at the tip of the cilia membrane (**Movie 1**), we ruptured the cilia membrane and established ‘whole-cilium’ patch recording to observe an outwardly rectifying current (**Figure 4A**, **Figure 4**-**Source Data 1**).

**Figure 4.**
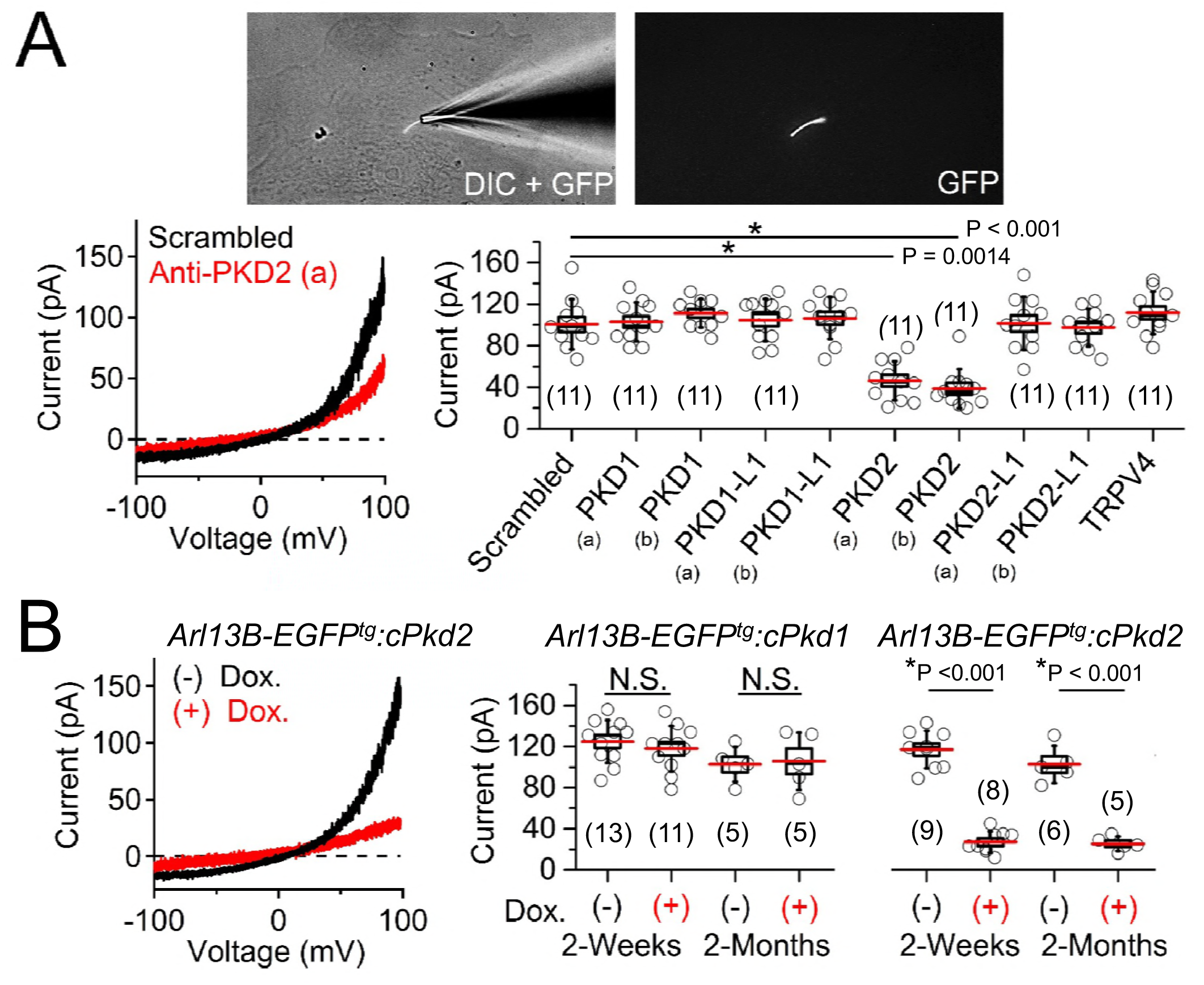
PKD2 is required for the ciliary ion channel conductance of the primary inner medullary collecting duct epithelial cells (pIMCDs). A) siRNA screen of potential I_cilia_ candidates. *Top*, light microscope image of a patched cilium. *Left*, Example ciliary currents measured from cells treated with either scrambled siRNA or one specific for PKD2. *Right*, Box (mean ± s.e.m.) and whisker (mean ± s.d.) plots of cilia total outward current (+100 mV) measured 48-72h after double-siRNA treatment. PKD1, PKD1-L1, PKD2 and PKD2-L1 mRNAs were targeted by two siRNAs specific for two different regions (a, b) of the target transcript (listed in **Table 1**). Averages are indicated by the red lines. Student’s *t*-test P values comparing treatment groups to scrambled siRNA. See “Figure 4-Source Data 1-ciliary currents for siRNA screen of TRP proteins in cilia”. B) Conditional knockout of the whole-cilia current. *Left*, exemplar cilia currents from pIMCD epithelial cells isolated from conditional PKD2 knockout (*Arl13B-EGFP^tg^:cPkd2*) transgenic mice. *Right*, box and whisker plots comparing the total outward cilia current (+100 mV) from control littermates and doxycycline-treated animals (*Arl13B-EGFP^tg^:cPkd1* and *Arl13B-EGFP^tg^:cPkd2*). Number of cilia is indicated by the italic number in parentheses for each group genotype and treatment group. Student’s *t*-test P values compare the outward cilia current from the untreated and doxycycline-treated animals. See “Figure 4-Source Data 2-whole ciliary currents in PKD1 or PKD2 knockout primary cells”.

**Table 1.** siRNAs used to screen for ciliary ion channel genes*

| siRNA gene target | ThermoFisher Silencer ID | siRNA location | % Knockdown efficiency |
| --- | --- | --- | --- |
| PKD1 (a) | 151949 | 999 | 75 ± 3 |
| PKD1 (b) | 151951 | 7303 | 73 ± 3 |
| PKD1-L1 (a) | n398242 | 1075 | 76 ± 2 |
| PKD1-L1 (b) | n398244 | 806 | 72 ± 3 |
| PKD2 (a) | 150154 | 1089 | 83 ± 2 |
| PKD2 (b) | 63551 | 488 | 81 ± 2 |
| PKD2-L1 (a) | 101318 | 350 | 84 ± 2 |
| PKD2-L1 (b) | 101422 | 830 | 82 ± 2 |
| TRPV4 | 182203 | 778 | 80 ± 2 |

\*Athanasia Spandidos, Xiaowei Wang, Huajun Wang, Stefan Dragnev, Tara Thurber and Brian Seed: A comprehensive collection of experimentally validated primers for Polymerase Chain Reaction quantitation of murine transcript abundance. *BMC Genomics* 2008, 9:633.

To determine the identity of this current, we treated cells with siRNA specific for members of the polycystin family and other localized putative cilia ion channel subunits(S. J. Kleene & Kleene, 2017; Kottgen et al., 2008; Yoder et al., 2002). We observed 53% and 61% attenuation of whole-cilium current from cells treated with two independent siRNAs targeted to PKD2 (**Figure 4A**, **Table 1**, **Figure 4-Source Data 1**). Importantly, we did not see any difference in currents when cells were treated with siRNAs targeting PKD1, PKD1-L1, PKD2-L1, and TRPV4, suggesting that none of these targets are essential subunits of the pIMCD ciliary current. To confirm these results, we measured ciliary current of pIMCD cells from *Arl13B-EGFP^tg^:cPkd2* mice at 2-week and 2-month after withdrawal of doxycycline treatment. As expected, the ciliary outwardly rectifying currents from the *Arl13B-EGFP^tg^:cPkd2* mice were reduced by 84% and 81% from 2-week and 2-month post-treatment groups compared to littermates not exposed to doxycycline (**Figure 4B**, **Figure 4-Source Data 2**). These results demonstrate that doxycycline-induced TetO-cre ablation of *Pkd2* stably abolishes the pIMCD ciliary current. In contrast, cilia records recorded from pIMCD cells isolated from doxycycline-treated *Arl13B-EGFP^tg^:cPkd1* mice do not have reduced current compared to cells from untreated animals from 2-week and 2-month post-treatment groups (**Figure 4B**, **Figure 4-Source Data 2**). From this data, we conclude that PKD2 is a subunit of a major ion current in renal tubule epithelial cilia and that the absence of PKD1 expression does not substantially alter the net PKD2 current in the cilium.

### Ciliary PKD2 preferentially conducts K^+^ and Na^+^ over Ca^2+^ ions

As discussed in the introduction, it is widely reported that calcium is a major charge carrier for PKD2 under physiological conditions. However, we find that the collecting duct epithelia cilia membrane is nearly 2.5 times more selective for potassium than sodium ions (relative permeability P_K_/P_Na_ = 2.4, **Figure 5-figure supplement 1A**), with little permeation by calcium (P_Ca_/P_Na_ = 0.06), barely different than presumably impermeant NMDG (P_NMDG_/P_Na_ = 0.05). Here, the relative permeability was estimated by the measured change in reversal potential when sodium was replaced by each test cation (**Table 3**). These data also demonstrate that PKD2’s selectivity is distinct from that previously reported for PKD1-L1/2-L1 recorded in the cilia of RPE and MEF cells, which is ~6x more selective for Ca^2+^ over Na^+^ and K^+^ (DeCaen et al., 2013).

**Table 2.**
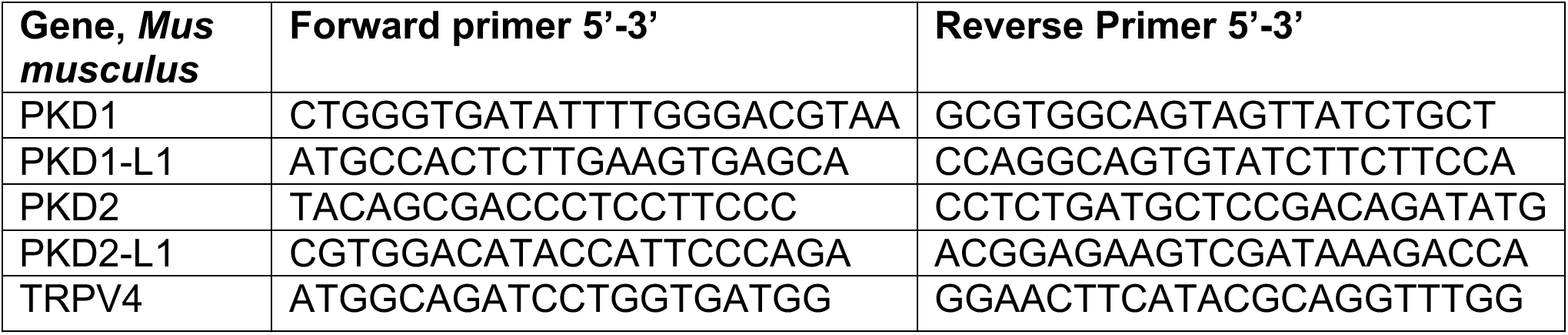
Primers used to detect gene expression using qPCR.

**Table 3.** Transmembrane potential reversal potentials (E_rev_) measured from pIMCD and HEK-293 PKD2-GFP cilia

| External Solution | pIMCD cilia |  | HEK293 PKD2-GFP cilia |  |
| --- | --- | --- | --- | --- |
| | $E_{rev}$ (mean ± s.d.) | $P_x/P_{Na}$ (mean ± s.d.) | $E_{rev}$ (mean ± s.d.) | $P_x/P_{Na}$ (mean ± s.d.) |
| NaCl | 0 | 1 | 0 | 1 |
| NaMES | -2 ± 3 mV | - | -1 ± 2 mV | - |
| KCl | 26 ± 3 mV | 2.7 ± 0.3 | 23 ± 4 mV | 2.4 ± 0.4 |
| CaCl<sub>2</sub> | -49 ± 4 mV | 0.08 ± 0.04 | -48 ± 3 mV | 0.09 ± 0.04 |
| NMDG | -57 ± 4 mV | 0.07 ± 0.05 | -58 ± 4 mV | 0.07 ± 0.05 |

We tested the effect of changing external calcium ([Ca^2+^] _ex_) while maintaining a constant level of Na^+^ (100 mM) on the magnitude on the inward ciliary current (**Figure 5-figure supplement 1B**). Here we observed that the inward current, presumably carried by Na^+^, was antagonized by [Ca^2+^] _ex_ (IC_50_ = 14 mM). This appears to be a consistent feature of PKD2 and mutated forms of the PKD2 channels when recorded from oocytes and reconstitution preparations(Arif Pavel et al., 2016; Cai et al., 2004; Koulen et al., 2002; Vassilev et al., 2001). To validate our findings from ciliary relative permeability, we also compared the single channel conductance of inward Na^+^, K^+^, and Ca^2+^ when they were exclusively present in the pipette (cilia-attached configuration). Of the three ions tested, K^+^ conducted through ciliary PKD2 channels with the greatest inward conductance (*γ*_K_ = 142 ± 6 pS), followed by sodium (*γ*_Na_ = 89 ± 4 pS) and calcium (γ_Ca_ = 4 ± 2 pS) (**Figure 5**). Importantly, inward calcium single channel currents were only observed when hyperpolarizing the cilia membrane potential (−140 mV to −200 mV) to increase the Ca^2+^ driving force (**Figure 5B**, **C**). Note that the outward conductance for all three conditions ranged between 98-103 pS, suggesting that the outward conductance is likely a mixture of Na^+^ and K^+^ exiting the cilium. The inward single channel open events were brief, usually lasting less than 0.5 ms (I_Na_ open time 0.4 ± 0.2 ms at −100 mV), whereas those measured at positive potentials opened for 190 times longer (I_Na_ open time 76 ± 29 ms at 100 mV). Thus, the order of ion conductance by PKD2 agrees with the relative permeability of the cilia membrane, from which we conclude that the major PKD2 conductance is monovalent, with very little inward Ca^2+^ flux.

**Figure 5.**
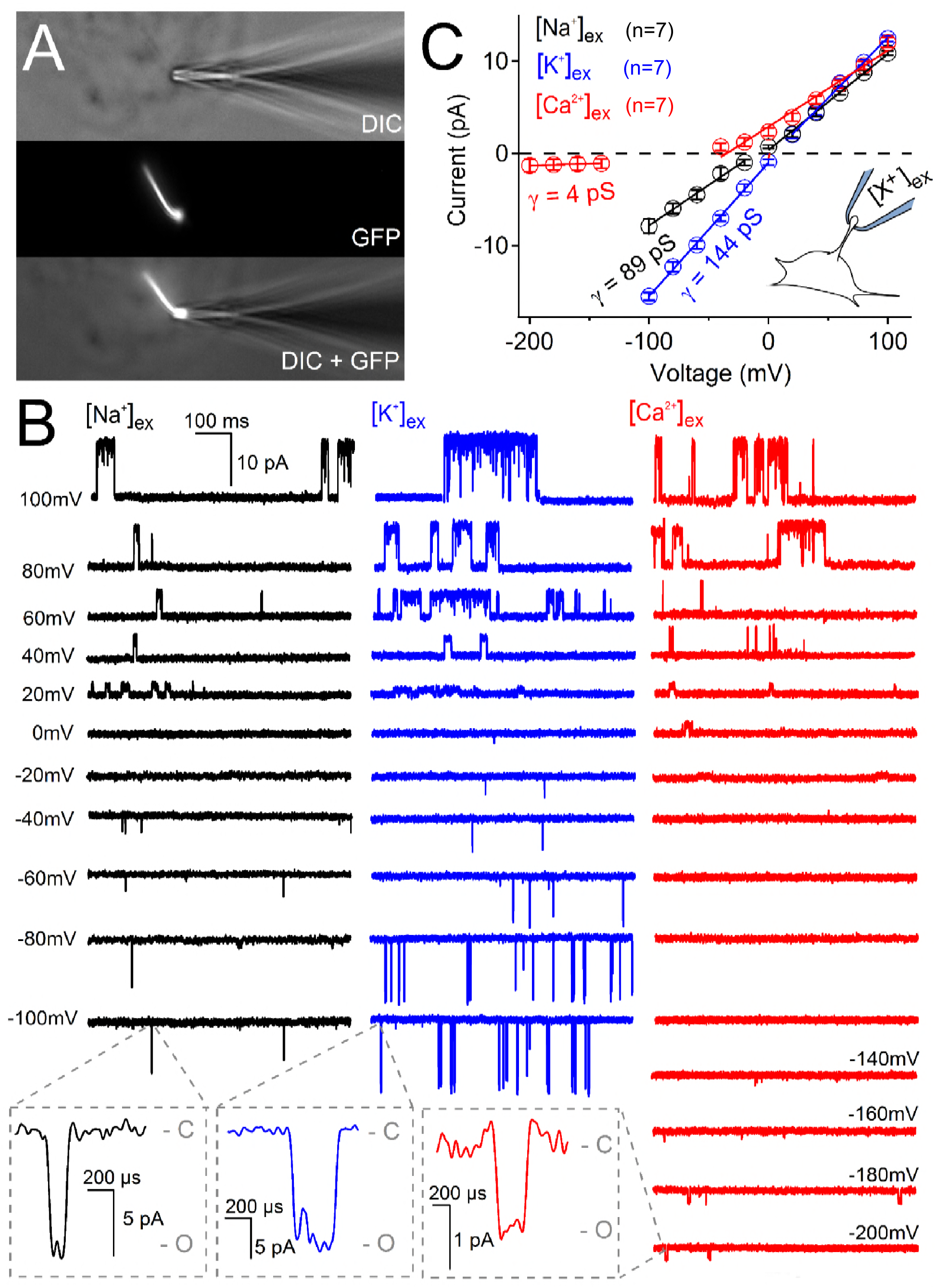
Ciliary PKD2 single channel currents conducted by sodium, potassium and calcium ions. A) Image of a cilia patched without breaking into the cilioplasm (‘on-cilium’ configuration). B) Exemplar currents recorded with the indicated cation (110 mM) in the patch electrode. Expanded time scales in the gray boxes show that inward single channel currents are brief, often opening (O) and closing (C) within 1 ms. C) Average single channel current amplitudes. Conductance (*γ*) estimated by fitting the average single channel currents to a linear equation. Note that the inward single channel events are very small (~0.8 pA at −200 mV) when Ca^2+^ is used as the charge carrier in the pipette (inset, patch diagram) whereas the outward current is much larger. Outward conductances of 90 pS, 99 pS and 117 pS, and inward conductances of 4 pS, 89 pS, and 144 pS were measured when the pipette contained Ca^2+^, Na^+^ and K^+^, respectively. *Inset*, a cartoon of the ‘on-cilium’ patch configuration where cations within the patch electrode ([X^+^]) are exclusively capable of conducting inward currents.

### PKD2 functions as a channel in primary cilia, but not in the plasma membrane

Since intracellular calcium has been reported to sensitize the ER-localized PKD2(Cai et al., 2004) and the polycystin-like channel (also called PKD-L or PKD2-L1)(DeCaen, Liu, Abiria, & Clapham, 2016), we examined this property in pIMCD ciliary PKD2 channels. Inside-out cilia membrane single channel activity can be compared to varying levels of intraciliary free calcium ([free Ca^2+^]_in_) (**Figure 6A**). Cilia patches most commonly had at least 3-4 active PKD2 channels per patch (**Figure 6B**), but some inside-out patches had only one PKD2 channel present. We used these rare patches to determine that 3 μM [free Ca^2+^]_in_ enhanced the open probability of the PKD2 current ~10-times (increasing P_o_ from 0.034 ± 0.02 to 0.36 ± 0.07) and the mean open time ~6-times (increasing from 37 ± 26 ms to 215 ± 40 ms) compared to standard cytoplasmic concentrations of 90 nM [free Ca^2+^]_in_ (**Figure 6B-D**). The half maximal [free Ca^2+^]_in_ enhancement of IMCD PKD2 open time was 1.3 μM.

**Figure 6.**
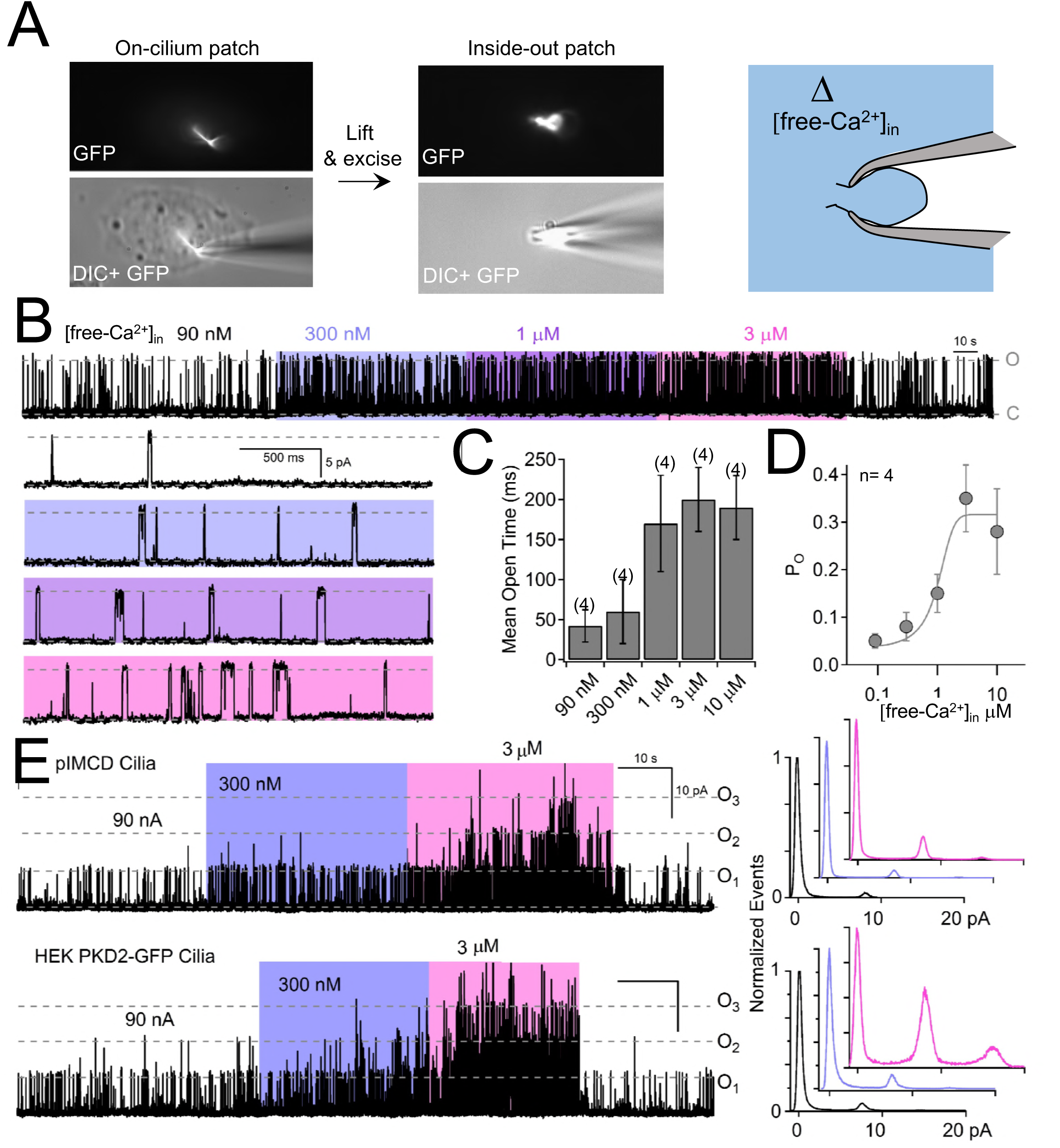
Internal [free-Ca^2+^] potentiates PKD2 channels. A) Images recorded while establishing an inside-out cilium patch. *Left*, a high-resistance seal is formed on the cilium; *Middle*, the electrode is then lifted, ripping the cilium from the cell body (see Methods); *Right*, cartoon depicting the inside of the cilium exposed to bath saline (blue) in which [Ca^2+^] can be adjusted. B) pIMCD PKD2 single channel events recorded in the inside-out configuration. The membrane potential was held at +100 mV in symmetrical [Na^+^] while the internal [Ca^2+^] was altered for 40-60 s intervals. C) The average open time duration relative to internal [Ca^2+^] (mean ± s.d.). D) Average open probability as a function of internal [Ca^2+^], fit to the Hill equation (described in Methods, mean ± s.d.). E) *Left*, Exemplar inside-out cilium patch records from pIMCD cilia and HEK-293 cilia with heterologously expressed PKD2 channels. *Right*, Current histograms capturing multiple open channel events under high internal [Ca^2+^] conditions. Currents were normalized to the closed (0 pA) state magnitude for each internal [Ca^2+^].

The above results confirm the location of functional ciliary PKD2 channels. Since the single channel recordings are made from the tips of cilia, PKD2 is present on the cilia membrane itself, not just the cilium/plasma membrane junction. However, there are reports that native PKD2 channel are constitutively active in the plasma membrane of immortalized cell lines derived from kidney epithelial cells (mIMCD-3 and Madin-Darby canine kidney, MDCK, cells)(Luo, Vassilev, Li, Kawanabe, & Zhou, 2003). To test this possibility, we voltage clamped the plasma membrane of pIMCD cells harvested from *Arl13B-EGFP^tg^*:*cPkd2* mice (**Figure 7A**; **Movie 2**). Here, we typically observed an outwardly rectifying Na^+^-permeant current, an inwardly rectifying K^+^ current, and an apparent voltage-gated Ca^2+^ current (**Figure 7B**). However, when PKD2 was conditionally ablated, there was no difference in plasma membrane currents densities (**Figure 7C**). PKD2 function has been implicated in calcium transients originating in the ER, plasma membrane, and cilium(R. Ma et al., 2005; Nauli et al., 2003; Q. Qian et al., 2003). However, the plasma membrane current-voltage relationship, inactivation kinetics and pharmacology are typical of L-type calcium currents and did not change when PKD2 was ablated in these cells (**Figure 7D, E**). Thus, our findings suggest that PKD2 does not constitute a significant portion of the plasma membrane current found in primary collecting duct epithelial cells and does not alter the native voltage-dependent calcium current.

**Figure 7.**
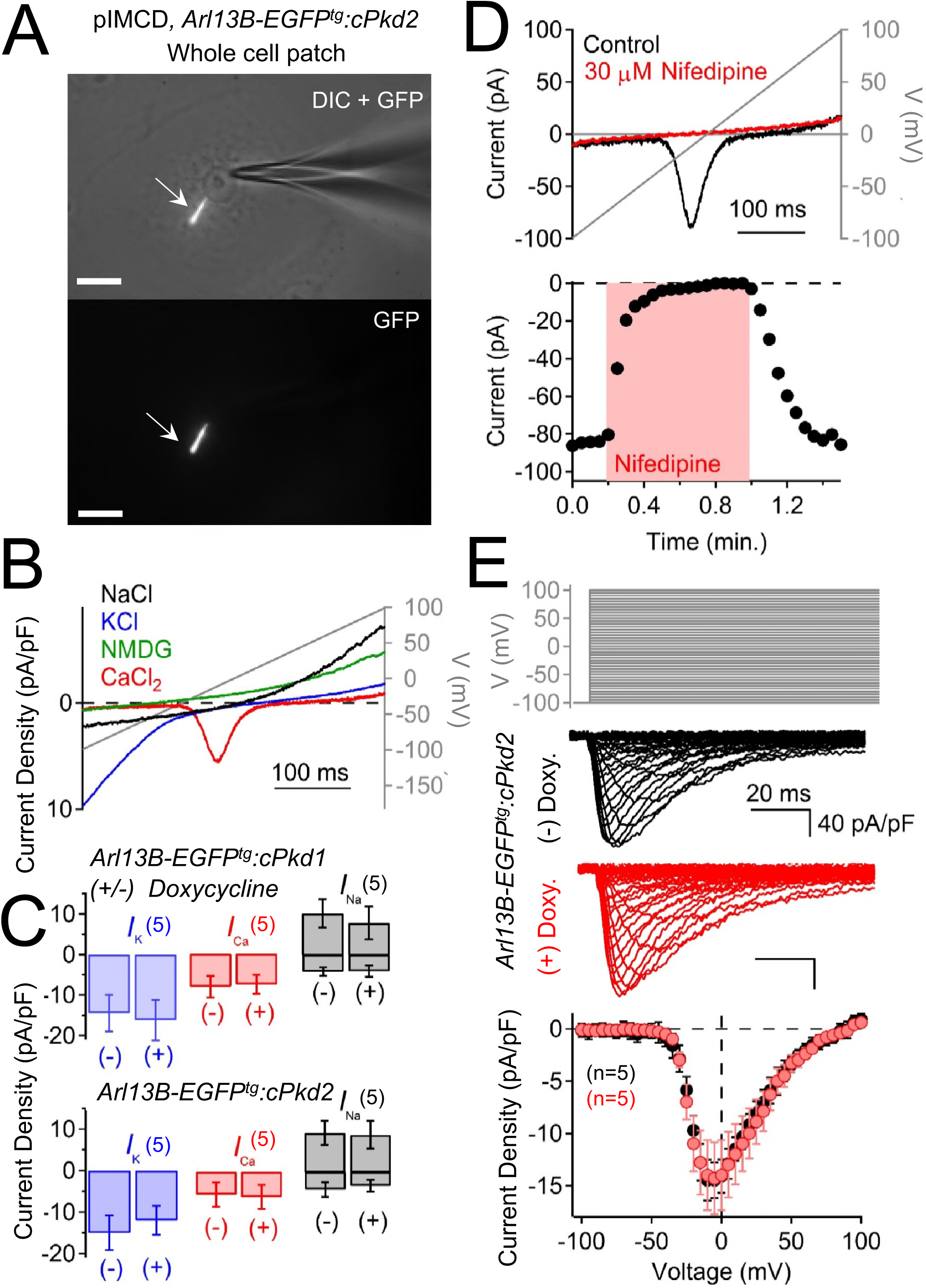
PKD2 does not generate a significant current in the plasma membrane of pIMCD cells. A) Images of a ciliated pIMCD epithelial cell patch-clamped on *the plasma membrane* (whole-cell mode). B) Example plasma membrane ionic currents activated by a voltage ramp (gray line) while changing the extracellular saline conditions. C) Average plasma membrane current density of *Arl13B-EGFP^tg^:cPkd2,* where animals treated with doxycycline in their drinking water was compared to untreated animals. Note that the average magnitudes of the plasma membrane currents were not significantly altered (mean ± s.e.m.). D) Pharmacological blockade of the voltage-gated calcium channel in the plasma membrane. Three independent experiments were performed. *Top*, Exemplar calcium currents blocked by nifedipine. *Bottom,* time course of block and recovery of Ca_V_ currents. E) Conditional PKD2 knockout does not alter the steady state voltage-gated calcium currents measured from the plasma membrane. *Top*, voltage protocol used to activate the calcium currents. *Middle*, exemplar leak subtracted voltage-gated calcium currents from *Arl13B-EGFP^tg^:cPkd2* animals, whose water was supplemented with doxycycline (red) and those whose water was untreated (black). Resulting plasma membrane average Ca_V_ densities compared from doxycycline-treated and control littermates.

### PKD2 is a ciliary channel but is not constitutively active on the plasma membrane in HEK-293 cells

Previously, our attempts to record heterologous PKD2 currents from the plasma membrane using transient transfection from multiple cell lines were unsuccessful. Thus, we generated two HEK-293 stable cell lines which overexpressed either human PKD2 with C-terminal (PKD2-GFP) or N-terminal GFP tagged (GFP-PKD2). In fixed preparations and live cells when viewed with confocal and standard fluorescence microscopy, GFP-PKD2 was intracellular, while PKD2-GFP also localized to primary cilia (**Figure 7-figure supplement 1A**; **Figure 7-figure supplement 2A, B**; **Movie 3**). The ciliary localization of PKD2-GFP in HEK-293 cells was confirmed in super-resolution images in which GFP co-localized with the known ciliary proteins, acetylated tubulin, and adenylyl cyclase (AC3) (**Movie 4; Movie 5,** respectively). However, we could not rule out PKD2 functioning in the plasma membrane of PKD2-GFP cells by fluorescence alone. To address this, we voltage clamped the HEK-293 PKD2-GFP whole-cell ion currents and found no difference when compared to parental HEK-293 cells (**Figure 7-figure supplement 1D**, surface area of cilia < 2% of total plasma membrane). Since native ciliary pIMCD PKD2 channels preferentially conduct K^+^, we compared the plasma membrane potassium current using steady-state voltage protocols. The resulting current-voltage relationship and block by 4-AP (4-aminopyridine, a K^+^ channel antagonist) suggests that the HEK-293 K^+^ current is conducted by native voltage-gated potassium channels (**Figure 7-figure supplement 1C**), commonly reported in these cells(Foley & Boccuzzi, 2010; Sands & Layton, 2014; Spandidos et al., 2008; Wilkes et al., 2017). Importantly, the kinetics and magnitude of the plasma membrane potassium current was not altered when PKD2-GFP was stably overexpressed (**Figure 7-figure supplement 1D**). Thus, overexpressed PKD2-GFP by itself does not appear to form a constitutively active or voltage-gated channel in the nonciliary plasma membrane.

However, when we patch clamped the ciliary membrane of the PKD2-GFP HEK-293 cells, a large outwardly rectifying single channel conductance was observed (**Figure 7-figure supplement 2C-E**). The PKD2-GFP HEK-293 ciliary inward conductance (*γ* = 90 ± 3 pS) was similar to the native PKD2 channels found in pIMCD cells (*γ* = 89 ± 4 pS) when sodium was used as a charge carrier. In the whole-cilium configuration, the cilia membrane primarily conducted K^+^ and Na^+^ with little permeation by Ca^2+^ (**Figure 7-figure supplement 3A, B**). To determine the identity of the outward rectifying current, we tested two siRNAs targeting to the overexpressed human PKD2 and observed a 29% and 32% decrease in outward current compared to scrambled controls (**Figure 7-figure supplement 3D**, **Figure 7-figure supplement 3D-Source Data 1**). To determine the localization of the channels mediating this current, we removed the cilia from the cell body by obtaining excised whole-cilium configuration recordings (**Figure 7-figure supplement 3C**). Excised patches from PKD2-GFP overexpressing HEK-293 cells retained an average of 44 ± 21% of the whole-cilium current. This percentage is highly variable since the amount of membrane preserved after excision varies between patches. Thus, it is unclear if the missing 56% of the ciliary current originated from the disconnected cilia membrane or near the cilia-plasma membrane junction. As found for the PKD2 channels in the pIMCD cilia, increasing internal calcium stimulated multiple open PKD2-GFP channels in the inside-out patch configuration of HEK-293 cilia (**Figure 6E**). Thus, ciliary PKD2 currents on PKD2-GFP overexpressing HEK-293 cells reproduces the ion selectivity and internal calcium sensitization as found in the native PKD2 channels of pIMCD cilia.

## DISCUSSION

### *Arl13B-EGFP^tg^:cPkd1* and *Arl13B-EGFP^tg^:cPkd2* as new mouse models for cilia visualization during ADPKD progression

We generated a mouse model in which expression of *Pkd1 or Pkd2* can be conditionally controlled in kidney tubule epithelia and confirmed cyst progression upon ablation of *Pkd1* or *Pkd2* in adults. Because the mice also expressed the *Arl13B-EGFP^tg^* transgene, we could compare the effect of polycystin ablation on cilia formation. We observed that conditional ablation of either gene did not block the formation of cilia in adult collecting ducts, consistent with findings in the embryonic node where constitutive global repression of *Pkd2* had no effect on cilia number(Field et al., 2011). Unexpectedly, we observed elongated and twisted cilia after *Pkd1* or *Pkd2* conditional ablation, a feature that became more pronounced as the ADPKD phenotype progressed. These data support the hypothesis that ciliary *Pkd1* and *Pkd2* are essential to the maintenance of renal tubular cell differentiation. However, we caution that the observed altered ciliary morphology may simply accompany cyst formation, and not necessarily be directly caused by the lack of these proteins within cilia. Because proteins change their subcellular localization during stages of the cell-cycle(Mollet et al., 2005; Morgan et al., 2002), it will be important to determine whether PKD1 or PKD2 localization changes during cell division when cilia are taken up, and if so, its biological significance.

### PKD2 is primarily a monovalent channel in the cilium

A commonly held hypothesis is that PKD1 and PKD2 form a calcium-selective channel directly involved in aberrant cytoplasmic calcium signaling(Pei, 2001). Previous work measuring reconstituted PKD2 channels from heterologous and native sources report conflicting voltage sensitivity and ion selectivity(Pablo, DeCaen, & Clapham, 2017). By directly measuring channels in primary cilia, we have shown that PKD2 is an essential subunit for the outwardly rectifying current. In primary cilia of primary cells native to kidney tubular epithelial cells, PKD2 ciliary current is relatively selective for monovalent cations (P_x_/P_Na_ = 2.4 and 1 for K^+^ and Na^+^), respectively, with comparatively little calcium permeation (Ca^2+^ ~ NMDG).

A recent report identified PKD2 as a large conducting ion channel (outward γ = 96 pS) of mIMCD- 3 cells(S. J. Kleene & Kleene, 2017). mIMCD-3 are immortalized epithelial cells derived from the terminal portion of the inner medullary collecting duct of SV40 transgenic mice and have similar morphology to the primary IMCD epithelial cells used in our study. We observed a similar outward conductance (*γ* = 98-103 pS) in the cilia of primary collecting duct epithelial cells directly harvested from adult mice. Like the results reported from the mIMCD-3 cell lines, unitary and whole-cilium currents from the pIMCD primary cells were abolished when PKD2 was knocked down with siRNAs or conditionally knocked-out in the whole animal. Both studies note the sensitization of the current by µM [Ca^2+^]. Based on these similarities, it is likely that we are describing the same ciliary PKD2 channel in these cell types. Both studies agree that the PKD2 cilia conductance is most selective for potassium but differ in estimates of sodium and calcium permeability (P_x_/P_K_ = 1; 0.14; 0.55 for K^+^, Na^+^ and Ca^2+^, respectively)(S. J. Kleene & Kleene, 2017) compared to P_X_/P_K_ = 1: 0.4: 0.025 for K^+^, Na^+^ and Ca^2+^, respectively, in this study. Apparent differences in the selectivity may arise from the differences in methodologies employed. In this manuscript, we record from primary tubule cells (pIMCD), not from immortalized cells (mIMCD-3), which are triploid. Second, external divalent contaminants were reduced by chelation in our solutions in which inward monovalent currents were measured. Finally, the method utilized by the Kleene group envelops the cilium within the recording pipette and forms a seal with the plasma membrane(N. K. Kleene & Kleene, 2012). Here, we are patching the cilium’s membrane exclusively (see **Movie 1**) and thus our membrane recordings are not affected by plasma membrane channels(DeCaen et al., 2013). It is important to note that neither we nor the Kleene group found inward calcium-mediated single channel events from potentials 0 mV to −100 mV (personal communication with the Kleene group). However, we were able to resolve unitary single channel events under non-physiological conditions in which the external calcium was high (110 mM) and cilia membrane potential was hyperpolarized (more negative than −120 mV). Thus, little calcium influx into the cilia would occur in the collecting duct since the resting cilia membrane potential is only −17 mV ^43^. Nonetheless, aberrant calcium signaling has been observed from cells expressing mutant PKD2 channels and interpreted as due to PKD2 function in ER calcium release(Cai et al., 2004; Celic et al., 2012). Since Ca^2+^ changes in the cytoplasm propagate into cilium(Delling et al., 2016), we cannot rule out the possibility that PKD2 in the ER alters [Ca^2+^]_cilium_. Also, although PKD2’s Ca^2+^ conductance is small, the cilium is a <1fL restricted space in which localized proteins might be influenced directly by occasional ciliary PKD2 channel Ca^2+^ flux. Most important, however, is the fact that the primary consequence of PKD2’s selectivity is to favor cilium depolarization and raise [Na^+^]_cilium_. If Na^+^/Ca^2+^ exchangers or Na^+^ -dependent kinases are found in cilia, PKD2 activity could underlie a slow, cumulative signal via calcium changes and/or kinase activity.

Renal epithelial cilia are exposed to urine, which contains varying ion concentrations as a function of position in the nephron. Human and murine distal collecting duct epithelial cells are exposed to high external concentrations of potassium (90-260 mM) and sodium (53-176 mM) ions, which contributes to the hyperosmolarity of urine (390-650 mOsm being considered normal, but can vary beyond this range depending on hydration state)(Callis, Vila, Carrera, & Nieto, 1999; Sands & Layton, 2014). Ciliary influx through PKD2, driven by these extreme extracellular concentrations of Na^+^ and K^+^ ions, may depolarize the plasma membrane sufficiently to activate voltage-gated calcium channels present in the plasma membrane. A recent computational study finds that opening of single ciliary PKD2-L1 channel (~ 150 pS) is sufficient to trigger action potentials in the soma of cerebrospinal fluid-contacting neurons at standard concentrations in blood plasma(Orts-Del'Immagine et al., 2016). Future electrophysiological studies using current clamp will determine whether the activation of ion channels in the ciliary membrane is sufficient to depolarize the plasma membrane of pIMCD cells, if there is any PKD1-dependent PKD2 function that has different ion selectivity or permeability compared to PKD1 independent function, or if there are more direct consequences of PKD2 expression for other ciliary compartment proteins.

### Ciliary PKD2 channels are activated by internal calcium

We have demonstrated that extraciliary Ca^2+^ can block monovalent conductance through PKD2, a common phenomenon in selective and non-selective cation channels, and responsible for the anomalous mole fraction effect (Eisenman, Latorre, & Miller, 1986; Friel & Tsien, 1989; Sauer, Zeng, Canty, Lam, & Jiang, 2013). This effect is likely due to a relatively higher affinity for Ca^2+^ ions in the pore (IC_50_ = 14 mM), thus blocking the channel to the inward passage of Na^+^ and K^+^ ions. Recently, PKD2 structures were captured in the ‘single’ and ‘multi-ion mode' states, in which 20 mM Ca^2+^ and 150 mM Na^+^ was present during protein purification(Grieben et al., 2017; Wilkes et al., 2017). Ca^2+^ bound at the entrance of the selectivity filter suggests Ca^2+^ blocks permeating Na^+^ ions in the multi-ion mode(Grieben et al., 2017; Wilkes et al., 2017). These studies provide a structural context for the anomalous mole fraction effects we observe in PKD2 currents from the pIMCD cilia and those reported by other groups measuring PKD2 channels from other preparations(Arif Pavel et al., 2016; Cai et al., 2004; Koulen et al., 2002; Vassilev et al., 2001). What, if any, effect might anomalous mole fraction effects have on the ciliary PKD2 channel? In contrast to the physiological, typically tightly-controlled interstitial [Ca^2+^] (~1.8 mM), urinary [Ca^2+^] in humans and mice is highly variable (5-20 mM) during normal diurnal activity(Foley & Boccuzzi, 2010). Urinary Ca^2+^ may have physiologically-relevant effects in dynamically limiting the PKD2 monovalent current through the cilium. When calcium-wasting occurs (~15 mM tubular [Ca^2+^]), more than half of Na^+^ and K^+^ influx through PKD2 would be antagonized (see **Figure 5-figure supplement 1**). Thus, ciliary PKD2 would be most active during low [Ca^2+^] in the tubule urine. On the other side of the cilia membrane, high internal [Ca^2+^] (EC_50_ = 1.2 μM) in the cilioplasm activates the PKD2 current by increasing its open probability ~25-fold. Internal Ca^2+^-dependent potentiation has been reported in PKD2 channels reconstituted from the ER into bilayers, but these channels inactivate at [free-Ca^2+^]_in_ concentrations >1 μM unless mutated at C-terminal phosphorylation sites(Cai et al., 2004). Previous measurements of resting [free-Ca^2+^]_in_ from RPE cilia (580 nM)(Delling et al., 2013) and from mouse embryonic node cilia (305 nM)(Delling et al., 2016) are 3-6 times higher than in the cell body. It is widely reported that in primary cilia, [free-Ca^2+^]_in_ increases to levels greater than 1 μM when mIMCD-3 cells (and other cell types) are exposed to flow or shear stress, based on the saturation of the fluorescence signal emitted by cilia-localized GCaMP and GECO calcium sensors(Delling et al., 2013; Su et al., 2013). Recent work has demonstrated that primary cilia are not Ca^2+^-responsive mechanosensors themselves; rather, mechanically-induced calcium waves are initiated from other locations to raise ciliary [Ca^2+^](Delling et al., 2016). Thus, increasing cytoplasmic [free-Ca^2+^] by mechanical or other stimuli, may increase cilioplasmic calcium and potentiate ciliary PKD2 channel activity.

### Polycystins PKD2 and PKD2-L1, independently form ion channels in primary cilia in disparate tissues

Previously we characterized the ciliary current from retina pigmented epithelial cells (RPE) and mouse embryonic fibroblasts (MEF) and demonstrated that they require the polycystins family, PKD1-L1 and PKD2-L1, based on attenuation of the cilia current by siRNA and genetic ablation of these two genes(DeCaen et al., 2013). Using similar methodology, including cilia electrophysiology from primary collecting duct cells, we have determined that PKD2 is at least a component of I_cilia_ from these cells. Thus, PKD2 and PKD2-L1 may be ciliary ion channels inhabiting distinct cellular tissues. Although the single channel conductance and sensitization of the RPE and MEF PKD1-L1/2-L1 cilia channel (Inward *γ*_Na_ = 80 ± 3 pS)(DeCaen et al., 2013) is similar to the pIMCD cilia PKD2 channel (Inward *γ*_Na_ = 89 ± 4 pS), their Ca^2+^ selectivity is distinct. Here we showed that the ciliary pIMCD PKD2 channel is blocked by external Ca^2+^ but is sensitized by high (EC_50_ = 1.2 μM) internal Ca^2+^, ~10 times the typical resting cytoplasmic concentration. Heterologous PKD2-L1 channels are also sensitized by increases in [Ca^2+^]_in_ (although the sensitivity range has not been determined) based on cytoplasmic Ca^2+^ uncaging studies and expected Ca^2+^ accumulation in whole cell experiments(DeCaen et al., 2016). However, a key difference between pIMCD (PKD2) and RPE (PKD2-L1) ciliary channels are that PKD2-L1 containing channels preferentially conduct Ca^2+^ (P_Ca_/_Na_ = 6-19)(DeCaen et al., 2013; DeCaen et al., 2016) over monovalent ions. Mutagenesis studies of heterologous PKD2-L1 channels has demonstrated that Ca^2+^ permeation is at least partly due to an additional glutamate residue (D525) on the external side of the selectivity filter, not present in PKD2(DeCaen et al., 2016).

Soon after these studies, PKD2 core structures were solved using single-particle electron cryo-microscopy in which the PKD2 channels formed a homotetrameric structure(Grieben et al., 2017; Shen et al., 2016), independent of coiled-coil domains (originally posited to form an interaction motif with PKD1 subunits(F. Qian et al., 1997)). As reported in Shen et al.(Shen et al., 2016), filter replacement of the PKD2-L1 channel with that of PKD2 conferred monovalent selectivity to the otherwise Ca^2+^-permeant PKD2-L1 channel. However, it is important to note that the native ciliary PKD2 channel’s ion selectivity, as described here, is not completely recapitulated in the PKD2 filter chimera, where single channel Ca^2+^ conductance was ~27-fold smaller than K^+^ (*γ*_Ca_ = 8 ± 2 pS, twice as large as the native cilia PKD2 channels *γ*_Ca_ = 4 ± 1 pS). Nonetheless, the native PKD2 cilia channel and the PKD2 filter chimera share similar ion selectivity, where K^+^ is favored over Na^+^ and Ca^2+^ as reflected in the magnitudes of single channel conductance (pIMCD cilia *γ*_K_ = 142 ± 6 pS, *γ*_Na_ = 89 ± 4 pS; PKD2 chimera *γ*_K_ = 218 ± 3 pS, *γ*_Na_ = 139 ± 3 pS) and relative permeability (P_x_/P_Na_ pIMCD cilia = 2.4: 1: 0.06 and PKD2 chimera = 2.2: 1: 0.5 for K^+^, Na^+^ and Ca^2+^ respectively). The physiological implications for the differential cilia expression of polycystins and attendant differences in ion selectivity is not clear.

### Loss of PKD1 does not alter PKD2 ciliary trafficking or PKD2-mediated ciliary currents

Based on the two-hit hypothesis of ADPKD, inherited haploinsufficiency of either PKD1 and PKD2 and a second acquired somatic mutation is required for disease progression. It has been reported that interaction between PKD1 and PKD2 is required for cell plasma membrane trafficking of the complex(Chapin, Rajendran, & Caplan, 2010; Gainullin, Hopp, Ward, Hommerding, & Harris, 2015; Hanaoka et al., 2000), but their interdependence for ciliary localization is still controversial. In some studies, PKD2 is absent from primary cilia without PKD1 or vice versa(Gainullin et al., 2015; H. Kim et al., 2014), while others showed PKD2 traffic to cilia is independent of PKD1(Cai et al., 2014; Geng et al., 2006; Hoffmeister et al., 2011). Our data support the view that PKD2 can traffic to primary cilia of pIMCD cells in the absence of PKD1. The disagreement of our data with some of previous reports may due to the different experimental systems. We employed isolated pIMCD cells with few or no passages, while others used cell lines (mIMCD3 cells, HEK-293 cells, LLC-PK1 cells, Renal Cortical Tubule Epithelial (RCTE) cell line) or cells overexpressing proteins. Another difference is that our mice express the cilia marker Arl13B-EGFP. There are no any other studies reporting this mouse model to date.

Based on biochemical data and initial whole cell electrophysiology, coexpression of PKD1 and PKD2 were shown to be necessary and sufficient to form heteromeric Ca^2+^-permeant channels in the plasma membrane(Hanaoka et al., 2000). However, we have noted a dearth of recordings in the literature, and of the few published, there are many inconsistencies. Based on the data here, and the selectivity measured from the PKD2 gating mutant (F604P)(Arif Pavel et al., 2016), PKD2 is primarily a monovalent channel with selectivity K^+^ > Na^+^, whose current is blocked by extracellular Ca^2+^. It resembles many TRP channels in being slightly outwardly rectifying under physiological conditions, with a large single channel conductance (~100 pS). PKD1 and PKD2 were believed to form a channel by association with their C-terminal coiled-coil domain(F. Qian et al., 1997). In contrast, recent structures of PKD2 homotetramers channels suggest that it can form a pore in the absence of the PKD2-coiled-coil domain and without PKD1 subunits(Grieben et al., 2017; Shen et al., 2016; Wilkes et al., 2017). Furthermore, removal of the conserved coiled-coil does not alter homomeric functional assembly or the related PKD2-L1 channel(DeCaen et al., 2016; Q. Li, Liu, Zhao, & Chen, 2002).

Our genomic PCR data (**Figure 1-figure supplement 2D**) cannot rule out the possibility that not all pIMCD cells completely lacked PKD1, although the more quantitative patch clamp results show no current differences (**Figure 4**) between doxycycline-treated and untreated groups. Our results cannot exclude the possibility that PKD1 may associate with PKD2 currents in the cilia or perhaps in the membrane of the endoplasmic reticulum or Golgi. It is also possible that PKD1 may still modulate ciliary currents, perhaps indirectly as a receptor for extracellular ligands, or through direct association. However, our findings demonstrate that PKD1 is not essential for basal activity of PKD2 in primary cilia of pIMCD cells. Further work will likely refine the mechanism of the two-hit model of ADPKD progression.

### PKD2 is not a constitutively active ion channel in the plasma membrane

In previous work, we established that PKD2-L1 can form a constitutively active ion channel in the plasma membrane and in primary cilia(DeCaen et al., 2013; DeCaen et al., 2016). We also reported that PKD2 did not appear to function on the plasma membrane, where HEK-293 and CHO cells transiently overexpressing PKD2, with or without PKD1, have the same level of plasma membrane cation currents observed in untransfected cells(DeCaen et al., 2013; Shen et al., 2016). In this manuscript, we present several lines of evidence suggesting that PKD2 does not constitutively function in the plasma membrane in kidney epithelial cells. First, conditional knockout of PKD2 does not alter the major cation currents found in the plasma membrane of primary collecting duct epithelial cells. Second, we did not observe differences in plasma membrane current measured from HEK-293 cells stably expressing PKD2-GFP in parental cells. While it is possible that PKD2 in the plasma membrane could be stimulated by a ligand such as Wnt3a, Wnt9b, and triptolide, we have not been able to reproduce activation of heterologous PKD2 with these reagents(S. Kim et al., 2016; Leuenroth et al., 2007; R. Ma et al., 2005). Functional PKD2, heterologously expressed in the plasma membrane of *Xenopus laevis* oocytes, required an F604P mutation near its intracellular gate(Arif Pavel et al., 2016), similar to mutations required for TRPML plasma membrane function(Grimm, Jors, Guo, Obukhov, & Heller, 2012; Xu, Delling, Li, Dong, & Clapham, 2007). The PKD2 F604P current was selective for potassium and sodium, but blocked by extracellular calcium and magnesium. Our attempts to record overexpressed PKD2 F604P membrane currents in mammalian cells have been unsuccessful(Shen et al., 2016), but such expression in *Xenopus* oocytes has been reproduced (unpublished data, courtesy of Michael Sanguinetti, Univ. Utah). What is clear is that *wt* PKD2 has no measurable constitutive activity in the plasma membrane in either mammalian or *Xenopus* expression systems.

### PKD2-GFP cilia recordings as a heterologous expression detection method

Other groups have observed epitope-labeled PKD1 and PKD2 in HEK-293 cell primary cilia(Gerdes et al., 2007; Lauth, Bergstrom, Shimokawa, & Toftgard, 2007). Here we used HEK-293 cilia electrophysiology as a more sensitive and specific functional assay. C-terminal GFP-tagged PKD2 enriches in primary cilium when stably expressed in HEK-293 cells. We also observed that N-terminal GFP-tagged PKD2 fails to localize to the cilium, suggesting that the sensor in this position interferes with trafficking. It is possible that the N-terminal GFP tag may interfere with the amino-terminal cilia-localization sequence (R_6_VXP)(Geng et al., 2006) and likewise, the C-terminal GFP tag may interfere with the ER retention sequence found in the C-terminus(Cai et al., 1999). However, since we have demonstrated that native (untagged) PKD2 is functionally expressed in cilia of collecting duct epithelia, the C-terminally-tagged PKD2-GFP over-expressed in HEK-293 cilia appears to properly localize. Like many overexpressed ion channels, we observed a high amount of GFP fluorescence from N- and C-terminally tagged PKD2 within intracellular compartments, such as the ER. PKD2 in the ER has been shown to be sensitized by cytoplasmic calcium, triggering Ca^2+^-induced Ca^2+^ release, possibly through direct interaction with IP3R channel(Koulen et al., 2002; Vassilev et al., 2001). However, we did not examine PKD2 function in the ER or how cilia PKD2 may alter intracellular store calcium release. Future work should determine if PKD2 channels function in ER membranes of native tissue and if differential localization confers unique channel properties. These findings present new avenues to study mutant forms of PKD2 which cause ADPKD. This method can possibly be used to determine which ADPKD forms of PKD2 are gain-of-function or loss-of-function, and perhaps alter channel trafficking to cilia. Ultimately, outcomes from these studies could form a rational basis for PKD2 dysregulation in ADPKD and enhance our basic understanding of ciliary ion channel function in cell biology.

## MATERIALS AND METHODS

### Electrophysiology

Ciliary ion currents were recorded using borosilicate glass electrodes polished to resistances of 14-18 MΩ using the cilia patch method previously described(DeCaen et al., 2013). Holding potentials were −60 mV unless otherwise stated. The pipette standard internal solution (SIS) contained (in mM): 90 NaMES, 10 NaCl, 10 HEPES, 10 Na_4_-BAPTA (Glycine, N,N′-[1,2-ethanediylbis(oxy-2,1-phenylene)]bis[N-(carboxymethyl)]-,tetrasodium); pH was adjusted to 7.3 using NaOH. Free [Ca^2+^] was estimated using Maxchelator(Bers, Patton, & Nuccitelli, 1994) and adjusted to 100 nM by adding CaCl_2_. Standard bath solution (SBS) contained 140 NaCl, 10 HEPES, 1.8 CaCl_2_; pH 7.4. Unless otherwise stated, ‘whole cilia’ ionic currents were recorded in symmetrical [Na^+^]. All solutions were osmotically balanced to 295 (± 6) mOsm with mannitol. Data were collected using an Axopatch 200B patch clamp amplifier, Digidata 1440A, and pClamp 10 software. Whole-cilia and excised patch currents were digitized at 25 kHz and low-pass filtered at 10 kHz. To accurately measure membrane reversal potential, five current pulses from voltage ramps were averaged. Extra-membrane conditions were changed using a Warner Perfusion Fast-Step (SF-77B) system in which the patched cilia and electrode were held in the perfusate stream. Data were analyzed by Igor Pro 7.00 (Wavemetrics, Lake Oswego, OR). The reversal potential, E_rev_ was used to determine the relative permeability of K^+^, Na^+^ and NMDG (P_X_/P_Na_) using the following equation:

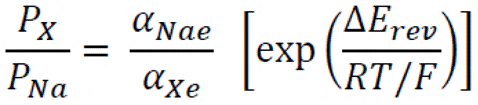

where E_rev_, *α*, R, T and F are the reversal potential, effective activity coefficients for the cations (i, internal and e, external), the universal gas constant, absolute temperature, and the Faraday constant, respectively. The effective activity coefficients (x) were calculated using the following equations:

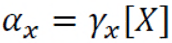

where *γ*_x_ is the activity coefficient and [X] is the concentration of the ion. For calculations of membrane permeability, activity coefficients (*γ*) were calculated using the Debye-Hückel equation: 0.79, 0.72 and 0.30 correspond to Na^+^, K^+^ and Ca^2+^, respectively. Since NMDG has no formal charge, it was assigned an activity coefficient of 1. To determine the relative permeability of divalent cations to Na^+^, the following equation was used:

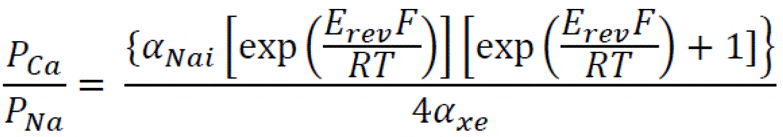

*E*_rev_ for each condition was corrected to the measured liquid junction potentials (-4.4 to 3.4 mV). The extracellular solution used 110 mM X-Cl, 10 HEPES and pH was adjusted with X-OH, where X corresponds to the cation tested. Monovalent-based extracellular solutions contained 0.1 mM EGTA to remove residual divalent cations. The NMDG-based extracellular solution contained: 110 NMDG; 10 HEPES; 0.1 EGTA and pH was adjusted with HCl. For ‘on-cilia’ single channel recording, the resting membrane potential was neutralized with a high K^+^ bath solution that contained: 110 KCl, 20 NaCl, 10 HEPES, 1.8 CaCl_2_, and adjusted to pH 7.4 using KOH. To test the inward single channel conductance, the intracellular pipette solution was replaced with one of the above extracellular solutions. For ‘inside-out’ cilia recordings, the pipette (extra-ciliary) solution contained SBS and the bath (intraciliary) solution contained 150 NaCl, 10 HEPES, 5 EGTA and free [Ca^2+^] adjusted by adding CaCl_2_. When a cilium was excised from the cell, the severed end of the cilium commonly re-seals itself, which isolates the intraciliary membrane from the bath and limits the effect of bath applied exchange on the inside of the cilium. To avoid this, excised cilia patches were briefly pressed against the surface of a bead made of Sylgard 184 (Dow Corning) to rupture the cilia at the opposing end. All electrophysiology reagents used were manufactured by either Sigma Aldrich or Life Technologies. The percent of inward I_Na_ block by external calcium was assessed by averaging the current at −100 mV before (I_control_) and after (I_Ca_) the addition of external and (I_Ca_ − I_control_/I_control_)×100. The potency of inward sodium current block was determined by fitting the percent inward current block and calcium concentration relationship to the Hill equation:

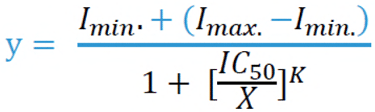

Where, I_min_. and I_max_. are the minimum and maximum response, IC_50_ is the half-maximum inhibition and K is the slope factor.

### Antibodies, reagents and mice

Mouse monoclonal antibody against GAPDH was from Proteintech. Rabbit polyclonal antibodies against acetylated tubulin(Lys40) was purchased from Cell Signaling Technology. Chicken polyclonal antibody against GFP was from Aves Labs. Doxycycline and Fluoshield^TM^ with 1,4-Diazabicyclo [2.2.2] octane were from Sigma-Aldrich; Hoechst 33342 was from Life Technologies. Rabbit polyclonal anti-mPKD2 (OSP00017W) and rabbit polyclonal anti-mAQP2 (PA5-3800) were from Thermo Fisher.

The *Pax8^rtTA^*; *TetO-cre*; *Pkd1^fl/fl^*; *Pax8^rtTA^*; *TetO-cre*; *Pkd2^fl/fl^* and *Arl-13B-EGFP^tg^* mice have been previously described(M. Ma et al., 2013)^,41^. *Pax8^rtTA^*; *TetO-cre* and *Pkd1^fl/fl^*; *Pax8^rtTA^*; *TetO-cre*; *Pkd2^fl/fl^* mice were obtained from the Somlo lab (Yale University). *Arl13B-EGFP^tg^:cPkd1 or Arl13B-EGFP^tg^:cPkd2* mice were generated by breeding *Pax8^rtTA^*; *TetO-cre*; *Pkd1^fl/fl^ or Pax8^rtTA^*; *TetO-cre*; *Pkd2^fl/fl^* with *Arl13B-EGFP^tg^* mice. The genotype was determined by PCR with the following primers. *Pax8^rtTA^* with PCR product ~600bp: IMR7385-CCATGTCTAGACTGGACAAGA; IMR7386 –CTCCAGGCCACATATGATTAG. *TetO-Cre* with PCR product ~800bp: TetO-CMV-5'- GCAGAGCTCGTTTAGTGAAC; Cre-R- TCGACCAGTTTAGTTACCC. *Pkd1^fl/fl^* with PCR product ~500bp (wild type ~300bp): ND1 Lox 5'-CACAACCACTTCCTGCTTGGTG; ND1 Lox 3'-CCAGCATTCTCGACCCACAAG. *Pkd2^fl/fl^* with PCR product ~450bp (wild type ~300bp): D2loxF1- GGGTGCTGAAGAGATGGTTC; D2loxR1- TCCACAAAAGCTGCAATGAA. *Arl13B-EGFP^tg^* with PCR product ~700bp (wild type ~400bp): 83940- TGCAACTCTATATTCAGACTACAG; 84608- GTGGACATAATGGTCCCATTAAGC; Transgene 2562- CATAGAAAAGCCTTGACTTGAGGTTAG. Mice were bred and housed in pathogen-free conditions with access to food and water in the Animal Care Facility. All experimental procedures were approved by the Boston Children’s Hospital Animal Care and Use Committee (IUACUC).

### Cell culture

Primary epithelial cells were cultured from dissected kidney collecting ducts of transgenic mice(Delling et al., 2016). Inner medullae were removed from the kidney and disassociated using a Dulbecco’s phosphate buffered solution (DPBS) containing 2 mg/ml collagenase A and 1 mg/ml hyaluronidase. After mechanical disassociation on ice, medullary cells were cultured in Dulbecco’s modified essential medium (DMEM) supplemented with 10% fetal bovine serum (FBS) and 100 units/ml penicillin/100 μg/ml streptomycin. Cilia were patched from cells after within 6 days after isolation and within one passage. siRNA knockdown efficiency was monitored for each experiment with a ‘scrambled’ silencer negative control 1 siRNA (Life Technologies). For generation of PKD2-GFP and GFP-PKD2 stable cell lines, C-terminal or N-terminal GFP-tagged PKD2 was generated by subcloning PKD2 cDNA into a modified pWPXLd vector. Puromycin (2 μg/ml) was used to screen stable cell lines. The PKD2 (NM_000297) human cDNA ORF Clone was purchased from Origene.

### Genomic PCR for pIMCD cells isolated from *Arl13B-EGFP^tg^:cPkd1* mice

Genomic DNA was extracted from the pIMCD cells isolated from *Arl13B-EGFP^tg^:cPkd1* mice (with and without doxycycline treatment). To genotype the *pkd1* alleles, primer 1 (P1), 5′-CCGCTGTGTCTCAGTGCCTG -3′, and P2, 5′-CAAGAGGGCTTTTCTTGCTG -3′, were used to detect the floxed allele (~400bp) and the *wt* allele (~200bp); while P1, 5 ′ - CCGCTGTGTCTCAGTGCCTG -3′, and P3, 5-ATTGATTCTGCTCCGAGAGG -3′, were used to detect the *Deletion* alleles (Deletion of exons 2-4 of *pkd1*) of *Del* (~650bp) and the non-Del (~1.65 kb).

### Immunocytochemistry, confocal microscopy and structured illumination microscopy (SIM)

Cells were fixed with 4% PFA, permeabilized with 0.2% Triton X-100, and blocked by 10% bovine serum albumin in PBS. Cells were labeled with the indicated antibody and secondary fluorescently-labeled IgG (Life Technologies) and Hoechst 33342. Confocal images were obtained using an inverted Olympus FV1000 with a 60x silicon oil immersion, 1.3 N.A. objective. Super resolution images using the SIM method where captured under 100x magnification using the Nikon Structured Illumination Super-Resolution Microscope (N-SIM) with piezo stepping. Confocal and SIM images were further processed with FIJI ImageJ (NIH).

### Immunohistochemistry and Cyst Parameters

*Arl13B-EGFP^tg^:Pax8^rtTA^;TetO-cre;Pkd1^fl/fl^* and *Arl13B-EGFP^tg^:Pax8^rtTA^;TetO-cre;Pkd2^fl/fl^* mice were induced with 2 mg/ml doxycycline in drinking water supplemented with 3% sucrose for 2 weeks from P28 to P42. Mice were anesthetized and perfused with 4% (wt/vol) paraformaldehyde at 8 weeks and 16 weeks after removal of doxycycline water. Kidneys were harvested and fixed with 4% paraformaldehyde at 4°C overnight, and embedded in paraffin. Sagittal kidney sections were stained with hematoxylin and eosin (H&E) and examined by light microscopy. All kidneys were photographed under the same magnification. ImageJ analysis software was used to calculate the cyst index (equal to the cumulative area of cysts within the total area of the kidney). For immunofluorescence of acetylated tubulin, a Leica VT1000S vibrating blade microtome was used for sectioning, kidney sections permeabilized with 0.5% TX100 / PBS pH 7.4 overnight, and blocked with BlockAidTM solution for 5-8 h. Sections were washed X3 in PBS, primary antibodies diluted in blocking solution, and sections incubated overnight at 4°C. After slides were washed X3 with PBS, goat anti-chicken/anti-rabbit fluorescent-labeled secondary antibodies were applied at room temperature overnight. Hoechst 33342 nuclear dye was incubated with sections for 1h. Sections were washed X3 with PBS, mounted in FluoshieldTM with 1,4-Diazabicyclo [2.2.2] octane and imaged with an inverted Olympus FV1000; silicon oil immersion 60x, 1.3 N.A. objective. Images were further processed and cilia length was measured using Fiji ImageJ (NIH).

### Inhibition and detection of transcripts

Approximately 200,000 primary cells were transfected with 100 pM of siRNAs (ThermoFisher) and 10 μl Lipofectamine^®^ RNAiMAX (Life Technologies) in a 9.5 cm^2^ dish. A list of the siRNAs is described in **Table 1**. At least 48h after transfection, half the cells were placed onto glass coverslips for electrophysiology, while the other half were lysed in TRIzol reagent (Ambion) for RNA extraction according to the manufacturer’s instructions. RNA was reverse transcribed using the SuperScript reverse transcription kit (ThermoFisher Scientific). Gene-specific primers were designed using Primerbank (http://pga.mgh.harvard.edu/primerbank/)(Spandidos et al., 2008). Transcripts were amplified by PCR and expression was visualized by agarose gel electrophoresis. Sequences for gene-specific primers are listed in **Table 2**.

## STATISTICAL ANALYSIS

Statistical comparisons were made using two-tailed Student's *t*-tests using OriginPro software (OriginLab, Northampton MA). Experimental values are reported as the mean ± s.e.m. unless otherwise stated. Differences in mean values were considered significant at P < 0.05.

## ACKNOWLEDGEMENTS

We would like to thank the Somlo lab (Yale University) for providing us with the kidney specific, doxycycline-dependent *Pax8^rtTA^; TetOcre*; *Pkd1^fl/fl^*(Shibazaki et al., 2008) and *Pax8^rtTA^; TetO-cre*; *Pkd2^fl/fl^*(M. Ma et al., 2013) knockout mice used to cross with the cilia reporter transgenic strain *Arl13B-EGFP*^*tg*^(DeCaen et al., 2013). We thank Nancy and Steve Kleene for the constructive discussions on the electrophysiology data. We thank the Northwestern Nikon Imaging Facility for use of their confocal and super resolution microscopes. We thank members of the Clapham and DeCaen labs for their productive discussions.

## AUTHOR CONTRIBUTIONS

X.L. generated the *Arl13B-EGFP^tg^:cPkd1*; *Arl13B-EGFP^tg^:cPkd2* animals and the PKD2 stably overexpressing cell lines; conducted and analyzed confocal and histology data.

P.G.D. performed confocal and electrophysiology experiments, and analyzed and interpreted the results.

X.L., P.G.D. and D.E.C. conceived and designed the experiments, analyzed and interpreted data.

T.V. conducted the Structured Illumination Microscopy experiments.

T.V., J.D. and S.S. assisted with the histology, generation and maintenance of the transgenic animals.

P.G.D. wrote the initial draft and X.L., T.V., P.G.D., and D.E.C. revised the manuscript.

## Figure Supplements Legends

**Figure 1-figure supplement 1.**
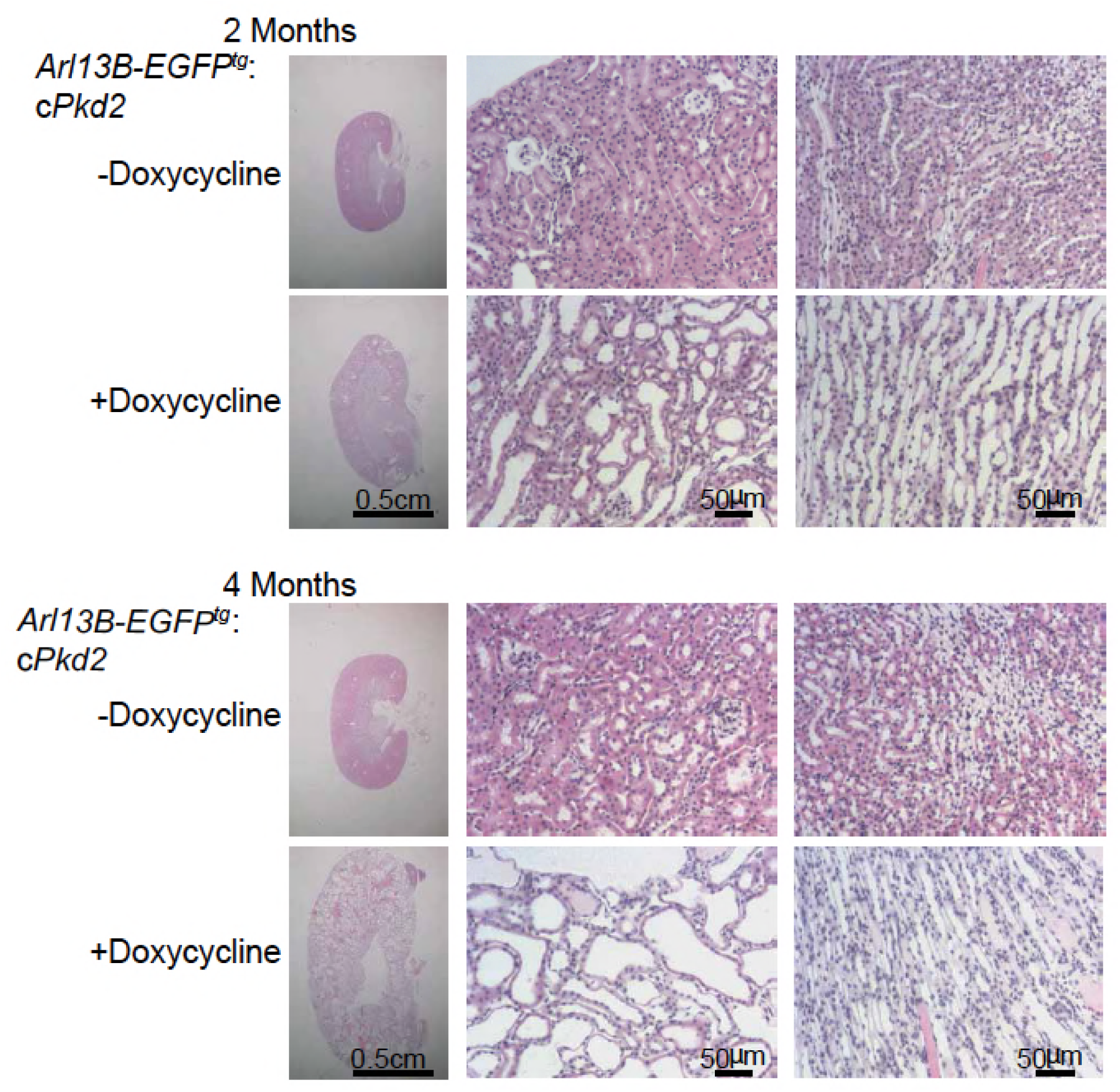
Progression of ADPKD after genetic ablation of PKD2. Representative images of H&E staining of kidneys from *Arl13B-EGFP^tg^:cPkd2* mice. Sagittal sections of kidneys (left, scale bar, 0.5 cm) and higher magnification (middle and right, scale bar, 50 μm) are shown at the indicated times. Three independent experiments were performed.

**Figure 1-figure supplement 2.**
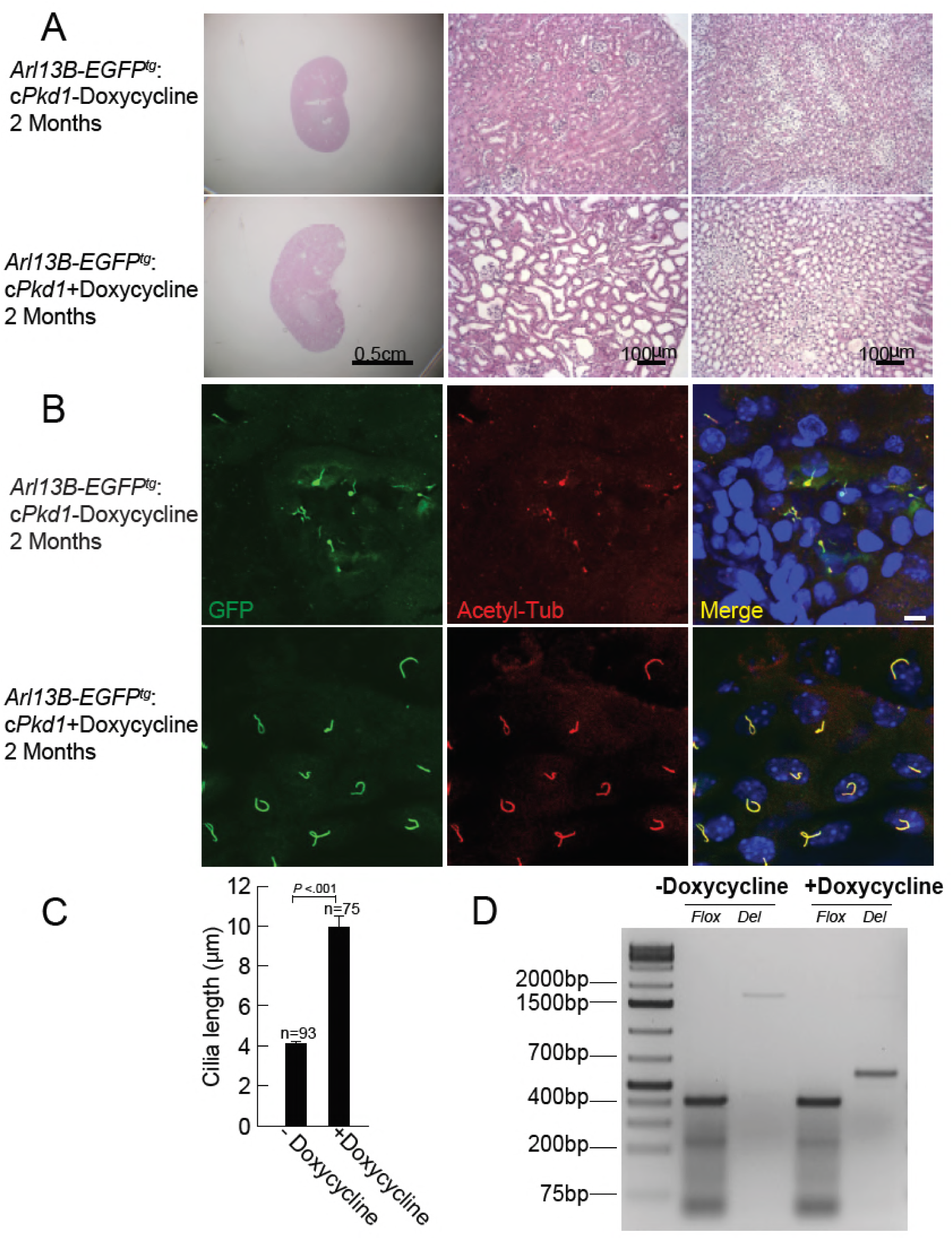
Progression of ADPKD after genetic ablation of PKD1. A) Representative images of H&E staining of kidneys from *Arl13B-EGFP^tg^:cPkd1* mice (2 months after doxycycline removal). Three independent experiments were performed. Sagittal sections of kidneys (left, scale bar, 0.5 cm) and higher magnification (middle and right, scale bar, 100 μm) are shown at the indicated times. B) Representative kidney sections from *Arl13B-EGFP^tg^:cPkd1* mice (2 months after doxycycline removal) were immuno-labeled with antibodies against EGFP and acetylated tubulin. Three independent experiments were performed. Scale bar, 5 μm. C) Cilia length increased with the progression of cyst formation from kidney sections of *Arl13B-EGFP^tg^:cPkd1* mice (2 months after doxycycline removal). D) Genomic PCR for pIMCD cells isolated from *Arl13B-EGFP^tg^:cPkd1* mice (2 weeks after doxycycline removal). Three independent experiments were performed. The PCR product for *floxed* allele, ~400bp; for *wt*, ~200bp; for the *Deletion* alleles (exons 2-4 of *Pkd1* deleted) of *Del, ~*650bp; for *non-Del,* ~1650bp.

**Figure 5-figure supplement 1.**
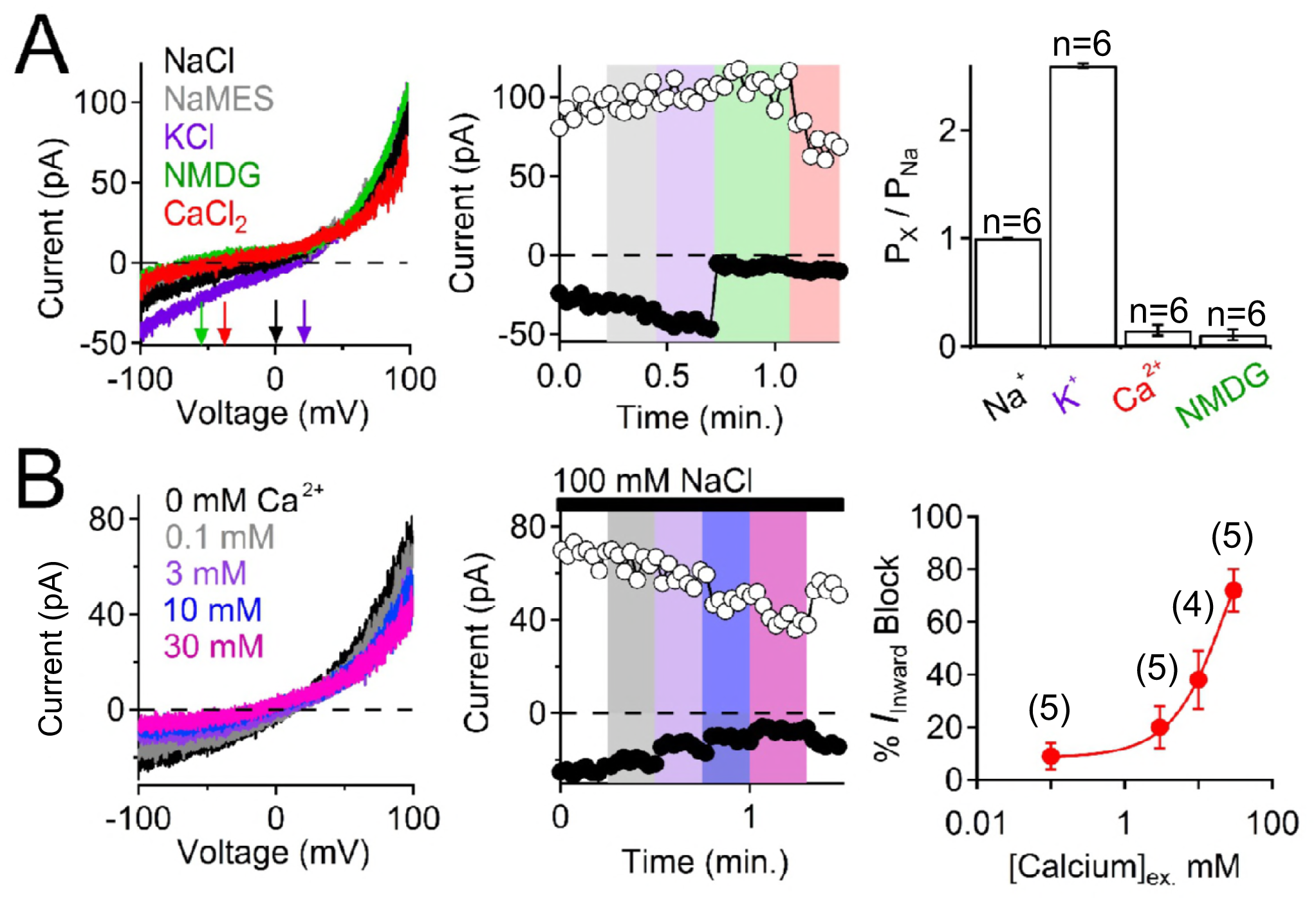
The pIMCD ciliary PKD2 cilia membrane is highly permeable to potassium and sodium monovalent cations. A) Selectivity of the ciliary PKD2 current. *Left*, Example cilia current traces measured under different external cationic bi-ionic conditions. *Middle*, Time course of the magnitudes of inward (-100 mV, black circles) and outward (100 mV, white circles) currents affected by changing the external solution. *Right*, Relative permeability of cations to Na^+^ through the PKD2 channel. B) Attenuation of inward PKD2 sodium current in response to external calcium ([Ca]_ex_). *Left*, the effect of external calcium on the outwardly rectifying PKD2 cilia current measured in symmetrical sodium. *Middle*, corresponding time course of the magnitudes of inward (-100 mV, black circles) and outward (100 mV, white circles) currents effected by increasing extracellular solution. *Right*, relationship between external calcium concentration and percent block of the inward sodium current (*I_inward_*) fit to the Hill equation. Number of cilia tested are indicated by the italic number in parentheses.

**Figure 7-figure supplement 1.**
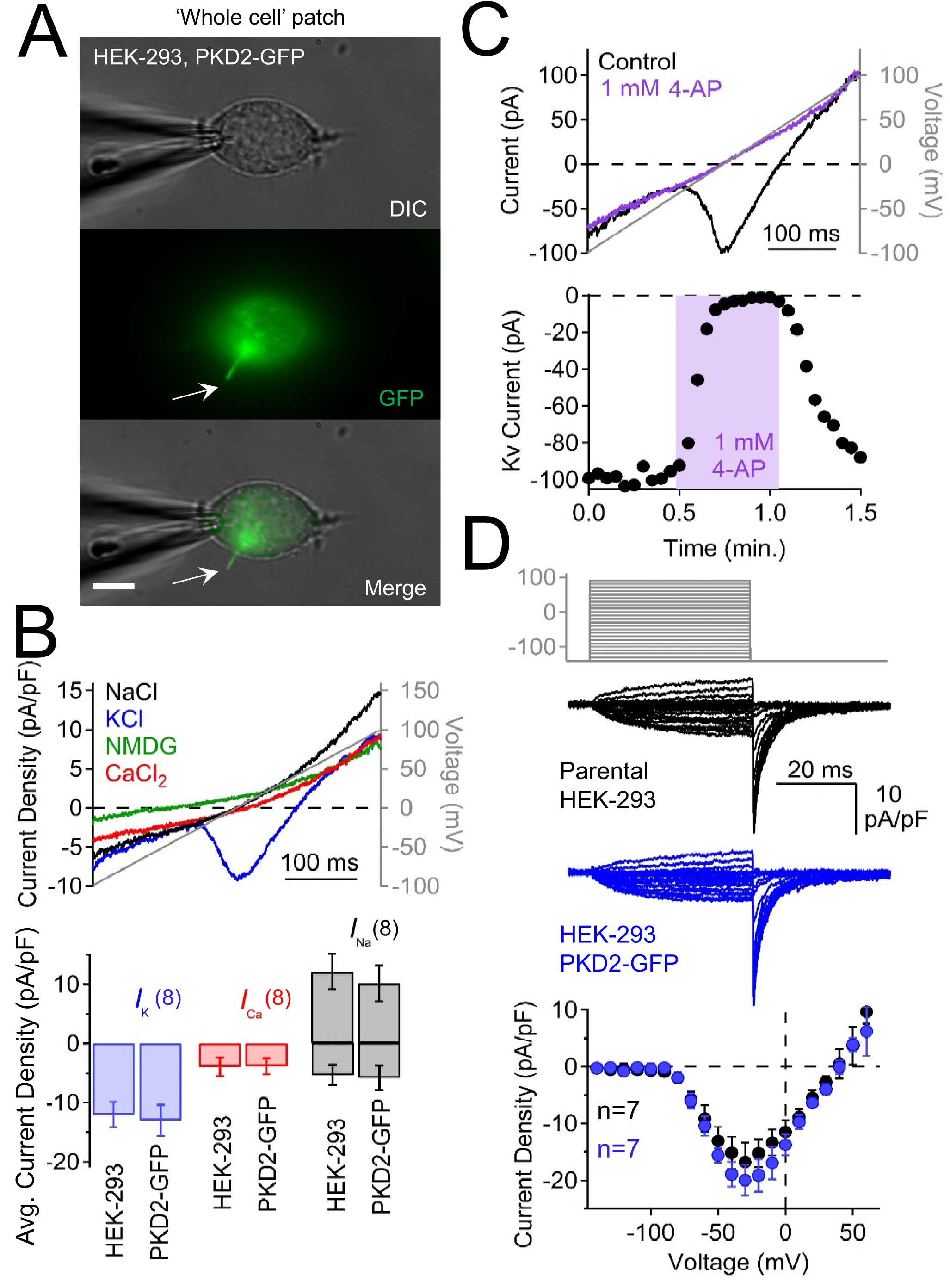
Characterization of plasma membrane currents measured from HEK-293 cells stably expressing PKD2-GFP. A) Image of a voltage-clamped HEK-293 cell stably expressing PKD2-GFP. Note that the GFP signal is present in the primary cilium (arrow) and that the patch electrode is sealed onto the plasma membrane. B) *Top*, resulting currents activated by a voltage ramp (gray line) under the indicated cationic conditions (in mM). Bottom, average current density for HEK-293 cells and those stably overexpressing PKD2-GFP. The current densities do not differ between the two cell types. C) Pharmacological blockade of the voltage-gated potassium current in the HEK-293 plasma membrane. Three independent experiments were performed. *Top*, Example voltage-gated potassium currents recorded before and after 4-aminopyridine (4-AP) exposure. *Bottom*, time course of block and recovery of potassium currents. D) Overexpressed PKD2 does not increase outward current (plasma membrane voltage-gated potassium current, K_v_). Steady state K_v_ currents measured from the plasma membrane. *Top*, voltage protocol used to activate K_v_ currents. *Middle*, exemplar leak-subtracted K_v_ currents from HEK-293 and those stably expressing PKD2-GFP. *Bottom*, K_v_ current density of HEK-293 and HEK-293 PKD2-GFP cells. Note that K_v_ current measured from HEK-293 cells is not different in HEK-293 cells overexpressing PKD2-GFP.

**Figure 7-figure supplement 2.**
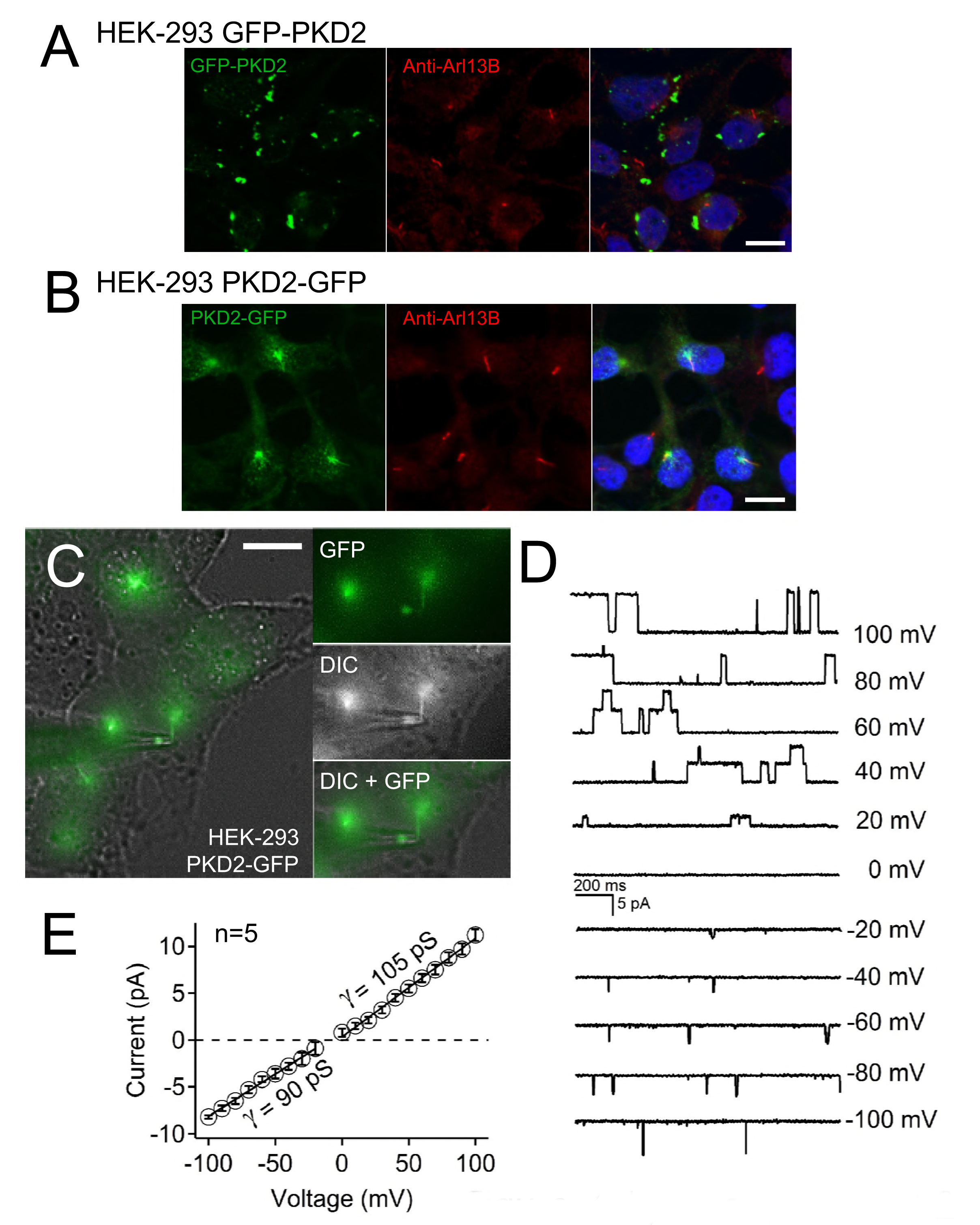
Single channel events recorded from cilium of PKD2-GFP overexpressing HEK-293 cells. A, B) Fixed HEK-293 cells stably expressing A) GFP-PKD2, or B) PKD2-GFP. Nuclei were stained with Hoechst 33342. Three independent experiments were performed. Scale bars = 10 μm. B) PKD2-GFP cells were immunolabeled with rabbit Arl13B primary antibody and detected by a secondary conjugated anti-rabbit Alexa 569 antibody (red). Three independent experiments were performed. Note that the Arl13B and PKD2-GFP signal colocalized to the cilium. C) Live cell image of an HEK-293 PKD2-GFP cilium patched in the on-cilium configuration. Note: the cilium’s tip is engulfed in the lumen of the electrode. D) Exemplar single channel currents recorded with a 100 mM NaCl-containing patch electrode. E) Resulting outward and inward conductances (*γ*) estimated by fitting the single channel currents to a linear equation.

**Figure 7-figure supplement 3.**
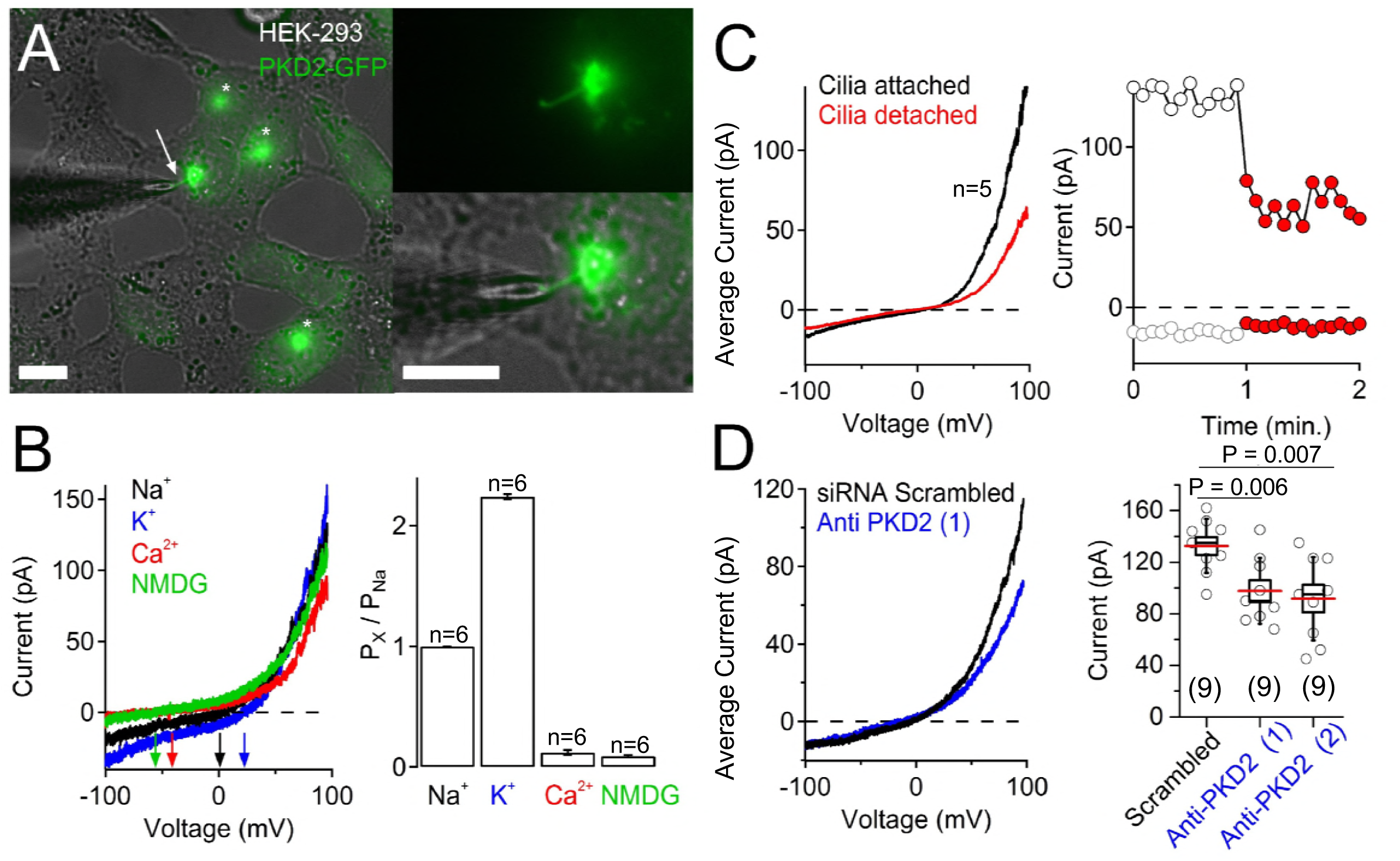
Overexpressed PKD2 forms an ion channel in the HEK-293 cilium. A) Image of a voltage-clamped cilium from HEK-293 cells stably expressing PKD2-GFP. Note that the GFP signal and/or cilia is not always present on HEK-293 cells. B) The selectivity of the ciliary current measured from HEK-293 PKD2-GFP cells. *Left*, Example ciliary current traces measured under different extracellular cationic bi-ionic conditions. *Right*, relative permeability of cations to Na^+^ through the PKD2 channel. C) The majority of the ciliary current is conducted through the ciliary membrane. *Right*, Average current traces from 5 voltage ramps acquired before and after the cilia is disconnected from the cell membrane. *Left*, time course of peak current amplitudes before (white circles) and after (red circles) cilia-cell separation. D) *Left*, average current traces from cells treated with scrambled or PKD2-mRNA-targeted siRNAs. *Right*, Box (mean ± s.e.m.) and whisker (mean ± s.d.) plots of ciliary total outward (+100 mV) current. Averages are indicated by the red lines. See “Figure7-supplement 3 Source Data 1-ciliary currents of cells treated with PKD2-mRNA-targeted siRNAs”. The GFP signal was noticeably affected in antisense-treated PKD2 cells and fewer cilia had as similarly intense fluorescence as the patched cilia (data not shown).

## Movie Legends

**Movie 1. Movie of the pIMCD Arl13B cilium patch configuration.**

The glass pipette patch electrode (*right*) is sealed onto a primary cilium above the pIMCD cell. The focal plane was moved along the z-axis (~9 µm) to visualize the cell and cilium. The electrode is moved along the y-axis while adjusting the focal plane to demonstrate that the patch electrode is sealed on the tip of the cilia membrane, not the cell membrane. The light source(s) are indicated in the lower left corner of the image: white light and 488 nm light to illuminate the specimen. Differential interference contrast (DIC) or fluorescent images were captured using a Hamamatsu Orca Flash CCD camera on an Olympus IX73 inverted microscope; 60x objective, 2x photomultiplier. Scale bar = 5 μm.

**Movie 2. Visualization of a plasma membrane patch from a ciliated pIMCD Arl13B-GFP cell.**

Same method as Movie 1. The cell membrane is being sealed by the patch electrode (*right*) and the primary cilium (*left*) is not in contact with the electrode. Scale bar = 10 µm.

**Movie 3. A plasma membrane patch established on a PKD2-GFP HEK-293 cell.**

Here the whole-cell configuration is formed by the patch electrode (*right*) on the cell body and the primary cilium (*lower right*) is not in contact with the electrode. The electrode is moved along the y-axis while adjusting the focal plane to demonstrate that the cell membrane is sealed on the patch electrode, not the cilia membrane. Scale bar = 5 μm.

**Movie 4. Ciliary co-localization of PKD2-GFP with immunolabeled acetylated tubulin.**

Paraformaldehyde-fixed HEK-293 cells stably expressing PKD2-GFP (*green*) were immunolabeled with anti-acetylated tubulin antibody (*red*). The images were captured using Structured Illumination Microscopy (SIM; Nikon N-SIM scope) and 3D images rendered using Imaris software (Oxford Instruments). Scale bar = 3 μm.

**Movie 5. Cilia co-localization of PKD2-GFP with immunolabeled adenylyl cyclase 3.**

Paraformaldehyde-fixed HEK-293 cells stably expressing PKD2-GFP (*green*) were immunolabeled with anti-adenylyl cyclase 3 antibody (*red*). The images were recorded using a Nikon N-SIM scope and 3D images rendered using Imaris software (Oxford Instruments). Scale bar = 3 μm.

